# Single-cell transcriptomic atlas of the human substantia nigra in Parkinson’s disease

**DOI:** 10.1101/2022.03.25.485846

**Authors:** Qian Wang, Minghui Wang, Insup Choi, Lap Ho, Kurt Farrell, Kristin G. Beaumont, Robert Sebra, John F Crary, David A. Davis, Xiaoyan Sun, Bin Zhang, Zhenyu Yue

## Abstract

Parkinson’s disease (PD) is a common and complex neurodegenerative disorder. Loss of neuromelanin-containing dopaminergic (DA) neurons in the substantia nigra (SN) is a hallmark of PD neuropathology; the etiology of PD remains unclear. Single-cell (-nucleus) RNA sequencing (sc or snRNAseq) has significantly advanced our understanding of neurodegenerative diseases including Alzheimer’s, but limited progress has been made in PD. Here we generated by far the largest snRNAseq data of high-quality 315,867 nuclei from the human SN including 9 healthy controls and 23 idiopathic PD cases across different Braak stages. Clustering analysis identified major brain cell types including DA neurons, excitatory neurons, inhibitory neurons, glial cells, endothelial, pericytes, fibroblast and T-cells in the human SN. By combining immunostaining and validating against the datasets from independent cohorts, we identified three molecularly distinct subtypes of DA-related neurons, including a *RIT2*-enriched population, in human aged SN. All DA neuron subtypes degenerated in PD, whereas the composition of non-neuronal cell clusters including major glial types showed little change. Our study delineated cell-type-specific PD-linked gene expression in the SN and their alterations in PD. Examination of cell-type-based transcriptomic changes suggests the complexity and diversity of molecular mechanisms of PD. Analysis of the remaining DA neurons of the three subtypes from PD demonstrated alterations of common gene sets associated with neuroprotection. Our findings highlight the heterogeneity of DA neurons in the human SN and suggest molecular basis for vulnerability and resilience of human DA neurons in PD. Our cohort thus provides a valuable resource for dissecting detailed mechanisms of DA neuron degeneration and identifying new neuroprotective strategies for PD.

## Introduction

The degeneration of dopaminergic (DA) neurons in the substantia nigra (SN) is a major pathological hallmark of Parkinson’s disease (PD). DA neurons regulate movement, learning, reward, and addiction. The loss of DA neurons in PD causes motor symptoms and can lead to psychiatric complications. The molecular mechanism that cause the loss of DA neurons in the SN remains poorly understood; several hypotheses, such as dopamine toxicity, iron burden, autonomous pace-making and axonal arborization, have been proposed to explain their vulnerability in PD^1^. However, detailed molecular and cellular dissection of human DA neurons is needed to understand the underpinning of DA neuron degeneration.

Identification of genetic variants and risk alleles of PD through genome wide association studies (GWAS) has begun to gain insight into the molecular mechanism of the disease^2^. However, the vast majority of PD cases have no known genetic cause, and their etiology remains unclear^3–5^. To dissect molecular mechanisms for midbrain DA neuron degeneration, post-GWAS research should investigate in what cell-type PD-linked genes or GWAS variants are actively expressed and how they are affected at the SN of PD brain.

Multiple studies in rodents have profiled DA neurons and demonstrated the heterogeneity of DA neurons in the midbrain. With single-cell transcriptomic analysis, they identified several molecularly distinct DA neuron subtypes in the midbrains, suggesting diverse functions and potentially differential vulnerability of different DA neuron types in PD^6–11^. By integration of GWAS and single-cell transcriptomic data from mouse brains, one study revealed an unexpected role of oligodendrocytes in PD progression^12^. While challenging due to sample scarcity and quality, few study has performed single-nucleus (sn)RNAseq in human postmortem midbrain and identified cell clusters representing DA neurons. Their results suggested an association of common risk for PD with DA neuron-specific expression^13, 14^. A recent study further examined PD midbrains by snRNAseq and reported a disease-specific DA neuron cluster and “pan-glial” activation ^15^. However, the subtypes or classification of DA neurons from the human SN remains uncharacterized^16^. Unlike mouse models, human DA neurons contain neuromelanin (dark pigment), which is biosynthesized from L-DOPA, a precursor of DA that increases in concentration during aging. While the loss of melanin-containing DA neurons in the SN has long been recognized in PD, it is underappreciated that a subpopulation of DA neurons in the SN of PD persisted through many years after the onset of motor symptoms, suggesting the resilience^17^. Indeed, how gene expressions and cellular functions are altered in the remaining DA neurons as well as other cell types in the SN from PD is largely unknown. Deciphering how the remaining DA neurons are adapted at the molecular level in the SN would provide valuable insight into the resilience or neuroprotective mechanism.

Here we reported snRNA-seq analysis of high-quality 315,867 nuclei from the postmortem SN of 23 PD and 9 non-PD controls (by far the largest) across various Braak stages (**Fig 1A).** The resulting transcriptomics atlas of the SN identified multiple cell clusters including DA neurons, excitatory neurons, inhibitory neurons, glial cells, endothelial, pericytes, fibroblast and T-cells. We identified three molecularly distinct DA related neuron subtypes that degenerate in PD. PD-linked genes show heterogeneous expression patterns in different cell types. Scrutinization of the remaining DA neurons of the three subtypes from PD reveals alterations of common gene sets associated with neuroprotection. Our transcriptomics data will be an important resource for future investigation into the detailed molecular mechanisms of the disease and discovery of therapeutic targets for PD.

**Fig. 1.**
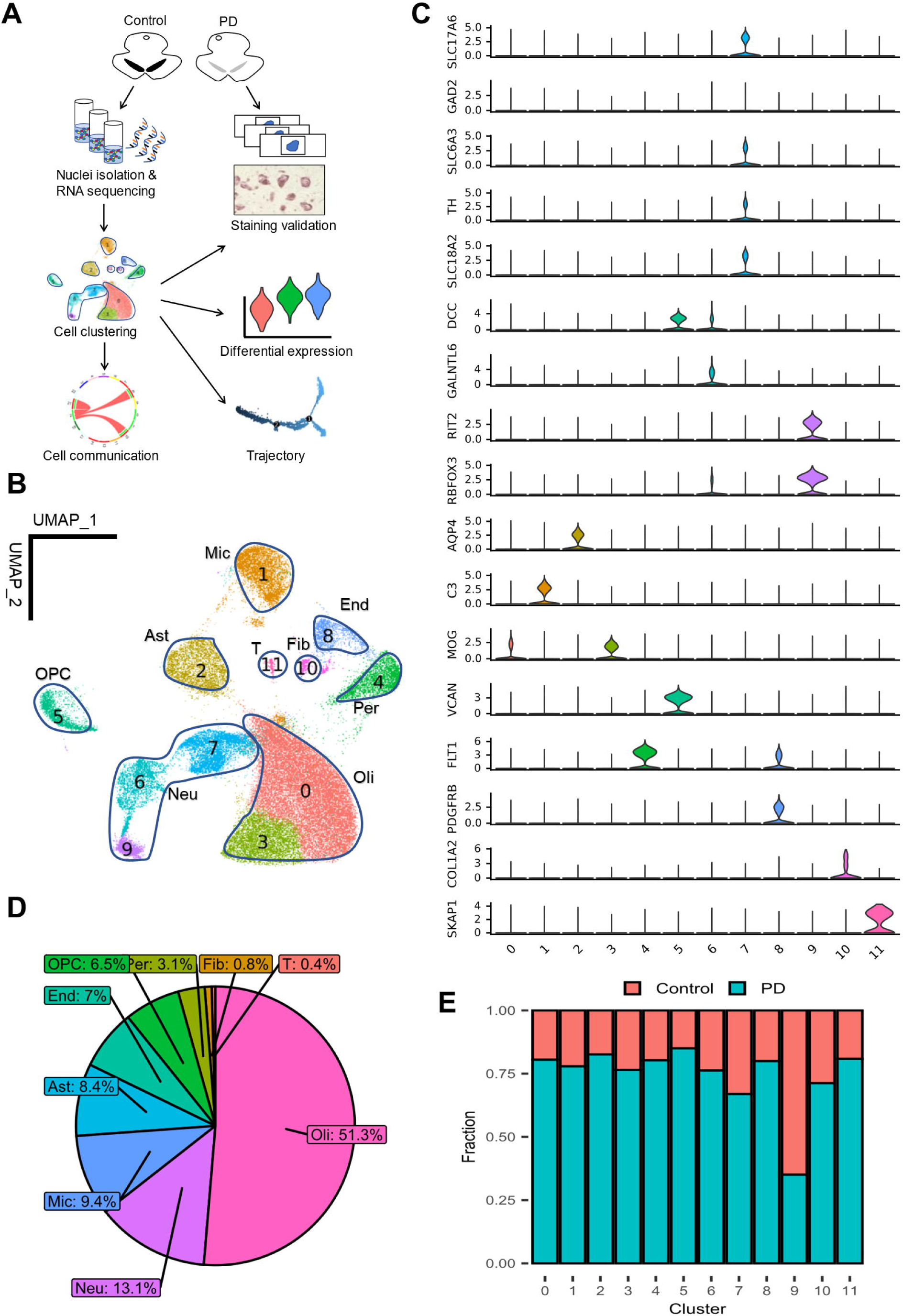
Cellular diversity in the substantia nigra from PD patients and control samples. **A**, Flow chart of the experimental procedure and data processing. Barcoded single-nucleus suspension was prepared using frozen SN samples from PD and the controls followed by RNA sequencing. Sequencing data were quality controlled and classified into cell clusters which were annotated with known cell type markers. Downstream analyses include cell composition changes, immunostaining, DEG identification, trajectory inference, and cell communication alterations. **B**, UMAP plot for cell clusters. Ast, astrocytes; Neu, neurons; Mic, microglia; Oli, oligodendrocytes; OPC, oligodendrocyte progenitor cells; End, endothelia; Fib, fibroblast-like cells; Per, pericytes; T, T cell. **C,** Expression pattern of known brain cell type marker genes in the control cells. **D**, Pie-chart for the fractions of major cell types in human SN. **E**, Sequenced cell distribution represented by disease status in each cluster.

## Results

### Cell-type composition and diversity in human substantial nigra

We have collected the SN from postmortem brains of 32 donors, including 23 idiopathic PD and 9 controls with an average age of 81, and processed them for snRNA-seq analysis (sequencing cohort, **Table 1** and **Table S1).** By using 10X Genomics Chromium Single Cell 3’ Solution, we obtained 457,453 droplet-based snRNA-seq profiles from these brains. Using a well-established Seurat-based data preprocessing^18^ and clustering analysis pipeline that includes quality control, data integration by Harmony^19^, cluster stability assessment, and doublets detection (detailed in Methods and **Fig. S1**), we obtained 315,867 high-quality nuclei and identified 12 cell clusters (c0 - c11) (**Fig. 1B**), which range in cell number from 134,011 (c0) to 1,384 (c11). To determine the cell-types of theses cell clusters, we employed two complementary strategies: (I) examining the expression pattern of known gene markers of major brain cell types, such as astrocytes (*AQP4*), neurons (*SLC17A6*, *GAD1*, and *RBFOX3*), microglia (*C3* and *CSF1R*), oligodendrocytes (*MOG*), oligodendrocyte progenitor cells (*VCAN*), endothelial (*FLT1*), and pericytes (*PDGFRB*) (**Fig. 1C**); (II) comparing *de novo* cluster-specific marker gene signatures (**Fig. S2A** and **Table S2**) with a large-scale collection of cell type markers curated from over 1,054 single-cell experiments^20^ (**Fig. S3**). Together, our annotation identified cell clusters that are categorized into 9 cell types, of which the identities and associated fractions are: oligodendrocytes (c0 and c3; 51.3%), neurons (c6, c7, and c9; 13.1%), microglia (c1; 9.4%), astrocytes (c2; 8.4%), endothelia (c4; 7.0%), oligodendrocyte progenitor cells (OPC) (c5; 6.5%), pericytes (c8; 3.1%), fibroblast-like cells (c10; 0.8%), and T cells (c11; 0.4%). Thus, the cell populations of the aged human SN are composed predominantly by oligodendrocytes, followed in descending order by neurons, microglia, astrocytes, endothelia, OPC, pericytes, fibroblast, and T cells (**Fig 1D**).

**Table 1.** Sequencing cohort demographic information. Data were represented by Mean±SD for age and PMI.

|  | Control (n = 9) | PD (n = 23) |
| --- | --- | --- |
| Age (years) | 83.8±8.3 | 78.8±7.6 |
| Sex: Male (%) | 4 (44) | 17 (73.9) |
| PMI (hrs) | 17.3±6.3 | 19.1±8.5 |
| Braak | NA | II:4; III:3; IV:5; VI:8; NA:3 |

### Identification of *RIT2*-enriched neuron subtype that degenerates in PD

When comparing the cell composition between the control and PD, we found that the fractions of the sequenced nuclei are nearly proportional to their sample size ratio (9 *vs* 23) in all clusters, except neuron cluster c9 (**Fig 1E**). While this observation demonstrates that the overall cell composition and most cell types are relatively intact in the SN of PD samples, c9 displayed a disproportionate distribution of the cell fractions between the control and PD, as compared to the rest of clusters (**Fig 1E**). To test the idea of neuron loss in the c9 cluster from PD samples, we calculated the odds ratio and performed immunohistochemistry (IHC) validation by using two separate cohorts (**Table S1).** In the first cohort, the control brains presented a significantly higher proportion of c9 neuron than PD brains (mean 3% vs 0.6%, overall odds ratio = 6.6, p = 0.0073 by Wilcoxon rank sum test of odds) (**Fig. 2A-B**). For the second cohort, we first stained the postmortem slices with an antibody against c9 neuron marker *RIT2* (**Fig. 1C**) and observed many *RIT2*+ neurons that also contain neuromelanin (NM) in the human SN *pars compacta* (SNpc) (**Fig. 2C**). While the NM+ DA neurons expressing TH are prevalent in control SNpc, we noticed a fraction of NM+ neurons lacking TH expression (**Fig. 2C-D**), consistent with a previous report that 7-30% of NM+ neurons are negative for TH staining^21^. We then perform co-staining with anti-RIT2 and anti-TH antibodies and found the partition of NM+ neurons into TH^High^ (54.25%) and RIT2^High^ (28.04%) subpopulations in control SNpc (**Fig. 2E**). Importantly, both TH^High^/NM+ and RIT2^High^/NM+ populations were markedly reduced in the SNpc of PD compared to that of the controls (**Fig. 2F**), corroborating the results from our snRNA-seq profiling. Thus, the observation shows the separation of RIT2^High^/NM+ neurons from TH^High^/NM+ neurons in control SNpc, while both degenerate in PD.

**Fig. 2.**
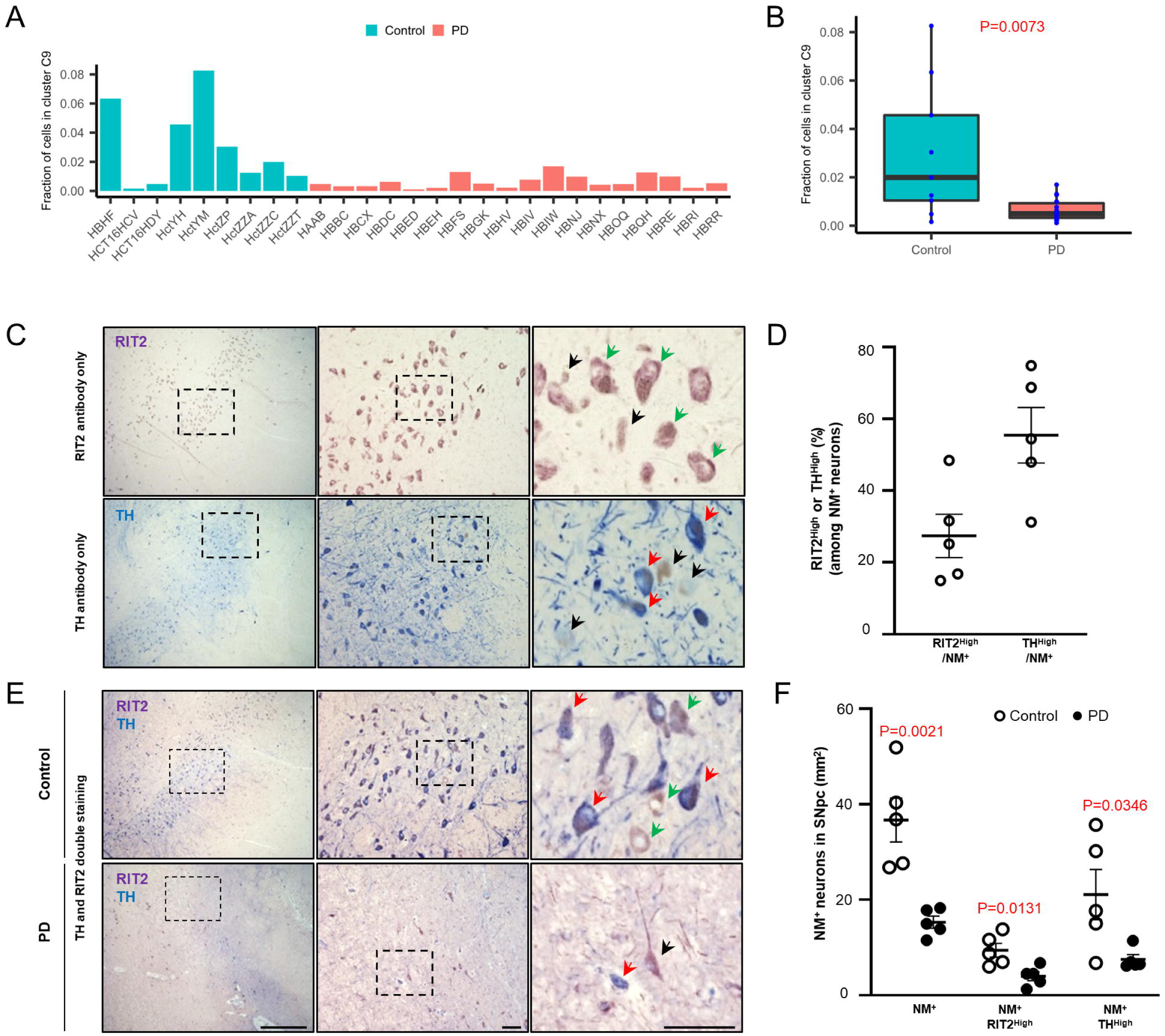
Loss of RIT2-enriched, neuromelanin-containing neurons in PD brains. **A**, Fraction of c9 neurons in one cohort containing PD and the controls. **B**, Distribution of the fraction of c9 neurons in PD and the controls shown in (**A)**. P value was computed by one-tailed Wilcoxon rank sum test. **C**, Paraffin-embedded sections containing human SNpc stained with antibodies against TH (blue) or RIT2 (purple) through chromogenic method. RIT2^High^ neuromelanin-containing neurons (NM^+^) (upper, green arrow), TH^High^ NM^+^ (lower, red arrow), and RIT2^-^ NM^+^ (black arrow) are indicated. **D**, Quantification of the ratio of TH^High^ or RIT2^High^ among NM^+^ neurons; **E**, Double staining with anti-TH and anti-RIT2 antibodies in the SNpc of PD and control brains. TH^High^ NM^+^ neurons (red arrow) and RIT2^High^ NM^+^ neurons (green arrow) are indicated. Scale bar, 1mm (left column); 100µm (middle and right column). **F**, Quantification of the number of NM^+^, TH^High^ NM^+^ and RIT2^High^ NM^+^ in the SNpc of the control (n=5) and PD (n=5). p-values were calculated by unpaired two-tailed Student’s *t* test.

*RIT2* encodes a neuronal GTPase and was shown enriched in DA neurons in the SN^22^. Interestingly, *RIT2* was previously identified as a PD susceptible gene^23^. We also noticed that a small fraction of c9 from the control (7.1%, 194 out of 2,741) express *TH*. To further validate the *RIT2*-enriched c9 neuron type in the human SN, we compared our results to published snRNAseq datasets from two independent cohorts containing only non-PD samples^14, 22^. We reprocessed their original datasets using our analytic pipeline to identify subtypes of the neuron clusters. In the dataset of *Agarwal et al* ^22^, we found a *RIT2*-enriched neuron sub-cluster (c6_0), which overlaps significantly with c9 in our study and is distinguished from their subcluster c6_1 enriched for typical DA markers such as *TH*, *SLC18A2,* and *SLC6A3* (**Fig. S4**). Note that the *RIT2*-enriched neuron subtype c6_0 from *Agarwal et al* express reduced levels *TH* and *SLC18A2* than c6_1. In the second dataset from *Welch et al*^14^, a *RIT2*-enriched neuron sub-cluster c4_5 overlaps significantly with c9 in our study and is separated from their c4_3, which is enriched for DA markers *TH*, *SLC18A2,* and *SLC6A3* (**Fig. S5**). Taken together, we conclude that *RIT2^High^*/NM+ (c9) represents a distinct subtype of DA neurons of human aged SN. c9 cluster is also enriched for additional markers, such as *RBFOX3, CADPS2, GRIK2, CDH18,* and *ZNF386D* (**Fig. S2A),** which distinguish itself from other cell clusters.

### Identification of subtypes of DA neurons that degenerate in PD

Neuron cluster c6 is enriched for marker genes *GALNTL6* and *DCC* (**Fig 1C**). *DCC* was shown to express at high levels in DA neurons in the ventral SN and modulate DA neuron viability^24^. To define the neuron type of c6, we performed sub-clustering analysis and obtained multiple subclusters of c6 (c6_0-5) **(Fig 3A-C)**. Focusing on subcluster c6_0-3 (c6_4 and c6_5 are excluded from further analysis due to small cell numbers from PD samples), we found that c6_2 is enriched for typical DA neuron marker genes such as *TH*, *SLC18A2,* and *SLC6A3.* Like c9, the fractions of c6_2 appear disproportionately distributed between the control and PD (49.7% from the control, overall odds ratio = 3.7, Wilcoxon p = 0.00096) (**Fig 3C),** suggesting the loss of c6_2 neurons in PD. By the similar approach, we obtained 4 subclusters of neuron cluster c7 (c7_0-3), and found a strong enrichment of *TH*, *SLC18A2,* and *SLC6A3* expression in subcluster c7_3 **(Fig 3D-F)**. Remarkably, c7_3 neuron number is also reduced in PD (60.8% from the control, overall odds ratio = 5.9, Wilcoxon p = 0.0041) **(Fig 3F)**. The expression patterns of distinct marker genes of DA neurons demonstrate the heterogeneity of DA neuron subtypes (**Fig 3G**) (**Fig S6A)**.

**Fig. 3.**
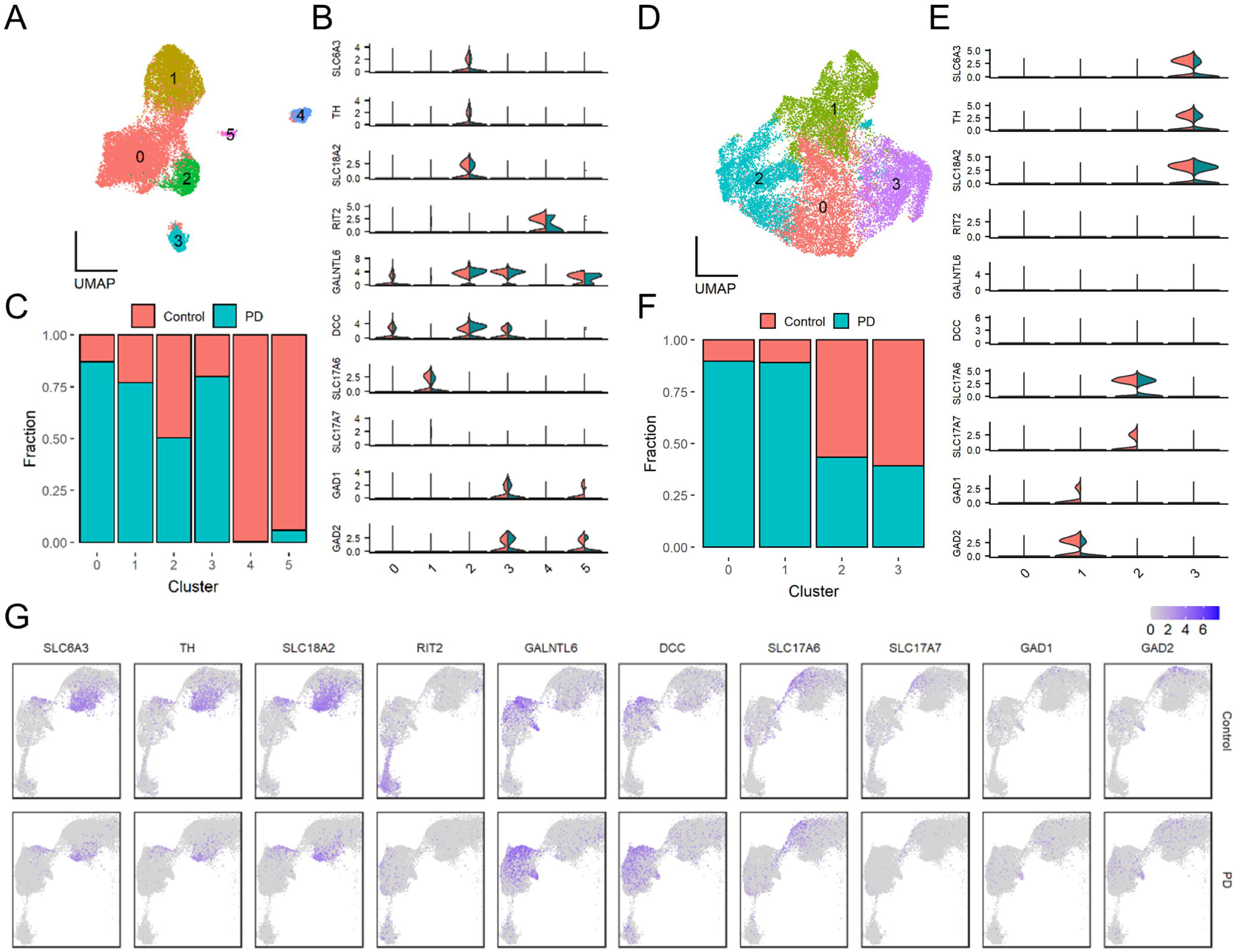
Sub-clustering analysis of clusters c6 and c7 and identification of subtypes of DA neurons. **A-F**, Sub-clustering analysis of cluster c6 (**A**-**C**) and c7 (**D**-**F**). **A** & **D**, UMAP plot for sub-clusters. **B** & **E**, Expression pattern of known brain cell type marker genes. **C** & **F**, Sequenced cell distribution represented by disease status in each sub-cluster. **G**, UMAP plots for the expression of selected marker genes in the neuronal clusters c6, c7 and c9.

Therefore, our results show three molecularly distinct clusters of DA neurons in the human SN: c6_2, c7_3, and c9, and they all degenerate in PD. Furthermore, while both c6_2, and c7_3 subclusters express high levels of *TH*, *SLC18A2,* and *SLC6A3* in the control and are considered typical DA neurons, they are distinguished by differential expression of different marker genes, such as high expression of *GALNTL6* and *DCC* in c6_2 not in c7_3. C9 are enriched for *RIT2* and *RBFOX3* expression and originated from DA neurons due to the presence of MN, thus representing a novel DA neuron subtype in human aged SN. The three DA neuron subtypes are distinguished by the differential expression of additional gene markers (**Fig. S2A, S6A)**.

Multiple studies have demonstrated the classification of DA neurons in mouse midbrain based on molecular profiling associated with the expression of distinct transcription factors (TF) confined in specific subtypes of DA neurons ^16^. By examining the TFs, many of which are critical for DA neuron specification or differentiation during development^25^, we found that the vast majority are expressed in a significantly low fractions of all cell clusters (including the three DA subclusters) in human aged SN from both the control and PD. *MYT1L* is an exception – it expressed in large fractions in the main neuron clusters (**Fig. S7**).

### Subtypes of glutamatergic neurons and GABAergic neurons in human SN

We observed that two neuron subclusters c6_1 and c7_2 are enriched for *SLC17A6,* which encodes vGlut2, suggesting that they are excitatory or glutamatergic neurons **(Fig 3B, E)**. In addition, c7_2 also expresses high levels of *SLC17A7* (vGlut1). To verify glutamatergic neurons in the SNpc, we stained the human SN slices with anti-vGlut2 antibody (**Fig. S2B**), and the results confirmed the existence of glutamatergic neurons in the SNpc, consistent with other reports^26, 27^. Interestingly, we also found a potential loss of c7_2 neuron in PD samples on average (odd ratio = 5), however, the Wilcoxon P value was insignificant at 0.799 **(Fig 3)**. Furthermore, we observed that C6_3 and c7_1 are both enriched for *GAD1* and *GAD2*, indicating that they are inhibitory GABAergic neurons in the human SN **(Fig 3B, E).** C6_3 differs from c7_1 by expressing high level marker genes *GALNTL6* and *DCC.* No apparent reduction of cell number is noted *in* C6_3 and c7_1 subclusters (**Fig 3 C, F**).

### Altered landscape of cell type-specific transcriptomics in the SN of PD

We next determined differentially expressed genes (DEGs) between PD and the controls in each cell cluster. DA neuron subcluster c9 and glutamatergic subcluster c7_2 presented with the highest numbers of DEGs, followed by endothelial cells (c4) and pericytes (c8) (**Fig. 4A** and **Table S3**). Functional enrichment analyses revealed upregulation of ribosomal genes and protein translation-related pathways in nearly all cell types (**Fig. 4B**). A broad increase of metallothionein family genes, such as *MT2A, MT1E*, and *MT3*, was also found in neuronal and non-neuronal clusters in the SN of PD (**Fig. 4C**). The metallothionein proteins are cysteine-rich and of low molecular weight. They bind heavy metal, and some appear to play a role in detoxification and cytoprotection^28^. In addition, a significant upregulation of several heat shock protein family members (e.g., *HSPB1, HSPH1, HSPA1, HSP90AA1*) as well as CRYAB, were observed in many cell types (including neuronal and non-neuronal clusters) in PD (**Fig. 4C**). *CRYAB,* encoding the alpha B subunit of Cystallin and a small chaperone protein associated with α-synuclein inclusion formation, was reported previously upregulated in the SN of PD^29, 30^. In contrast, vesicle trafficking, synaptic transmission and synapse-related genes were significantly downregulated in the neuronal clusters of PD brains (**Fig. 4B, 4C**).

**Fig. 4.**
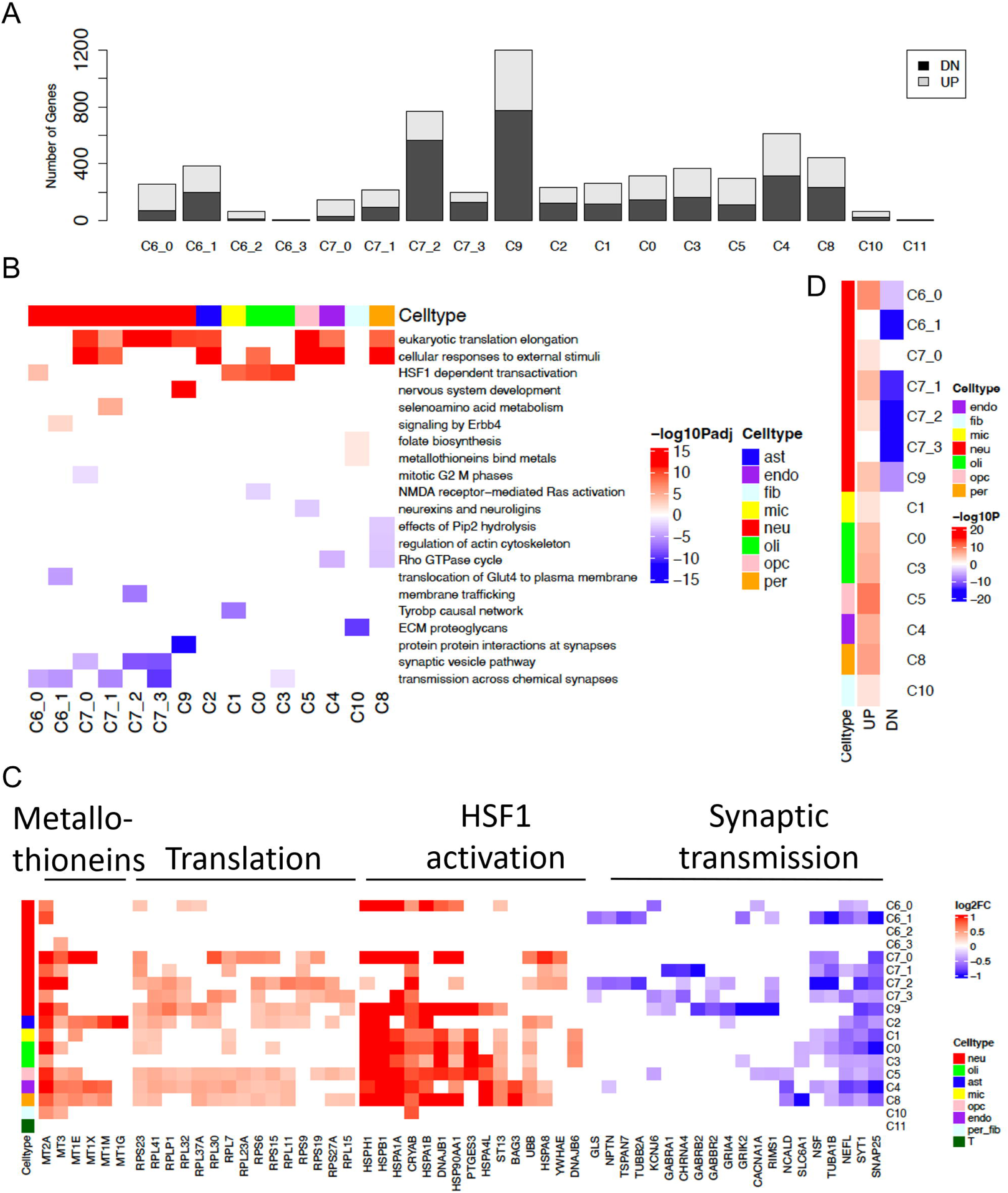
Cell-type-specific differential gene expression (DEG) between PD and the control. **A**, Number count of up- (UP) and down-regulated (DN) DEGs in each cluster. **B**, Heatmap of top canonical pathways enriched for up- and down-regulated genes in each cluster. **C**, Top DEGs involved in the indicated pathways in each cluster. **D**, Comparison of DEGs identified in bulk-tissue-based meta-analysis and snRNA-seq.

We noticed distinct patterns of DEGs among the three DA neuron subtypes in PD. c9 subtype had the greatest number of DEGs (1203), far more than c7_3 (200) and c6_2 (66). The most significant DEGs in c9 were associated with the down-regulation of synaptic protein interactions. In contrast, the DEGs of c7_3 showed the most significant upregulation of translation elongation and ribosomal proteins (**Fig 4B**). Moreover, c7_3 had significantly decreased expression of *SLC18A2, ALDH1A1, SLC6A3*, and *TH,* which are important for the regulation of dopamine release and dopaminergic neurogenesis. A few genes sharing the similar directions of changes (either up or down) among c9, c7_3 and c6_2 were also identified and they were involved in various cellular functions (**Fig. S6B-C**). Genes involved in single-stranded DNA sensing process such as *SSBP3* were commonly downregulated, while phospholipase C like 1 (*PLCL1*) and pseudogene *MTRNR2L1* were enhanced in all 3 DA clusters.

Finally, we observed impaired Tyrobp causal network in the microglia c1, indicating the immunosuppression in PD brains (**Fig 4B**). These cluster-specific PD DEGs were largely consistent with those identified in our previous bulk-tissue-based meta-analysis^31^, with the downregulated bulk-tissue-based DEGs most significantly enriched in neuronal clusters while others were found ubiquitously upregulated across various cell types (**Fig. 4D**).

To identify potential temporal changes of gene expression during disease progression, we separated the cells into three groups according to the Braak Staging of the donors: Control (Braak=0), early-stage (Braak 1-3), and late-stage (Braak 4-6) (**Table S4**). The genes were grouped into three major categories based on their expression patterns (details described in **Materials and Methods**): 1) Early and Sustained Responding Genes (ESRG); 2) “U-shaped” Responding Genes (URG) whose early response diminished over time; 3) Late Responding Genes (LRG). Each category was further divided into positive and negative responders. The positive ESRG were primarily involved in HSF1 activation in non-neuron clusters (c0∼c3, c5 and c8) and a small set of neuron clusters (c6_0 and c7_0), while in DA neuron subclusters c7_3 and c9, DA neurogenesis and synaptic homeostasis, respectively, were disrupted at the early stage and then suppressed over the disease course (**Fig. S8A**). Most positive URGs were found enriched in metal ion regulation and metabolism in non-neuronal clusters while a transient inactivation of NMDA-AMPK signaling and membrane trafficking was found in oligodendrocyte (c0) and microglial (c1) clusters (**Fig. S8B**). Various cellular pathways were activated in all cell clusters as LRGs, including the two vulnerable DA neuron subtypes (c7_3 and c9), where translation elongation, chaperone activity and GTPase cycle are altered (**Fig. S8C**). Moreover, Tyrobp causal network and microglial pathogen phagocytic pathway was interrupted at late stage (**Fig. S8C**). The results demonstrate divergent cellular stress responses in different cell types during disease progression.

### Effect of sex in differential gene expression of the SN in PD

Sex is a known risk factor for PD as men develop PD more frequently than women^32^. To investigate if there exists sex-specific gene dysregulation in PD in the SN, we conducted DEG analysis in each cluster for males and females separately (**Table S5**). Males and females shared most DEGs (656) in c9. (**Fig. S9A**). While sample size difference may partially explain the difference in the number of sex-specific DEGs between sexes (21 males vs 10 females and hence more male cells than female cells in all clusters except c9), there were more female specific DEGs than males in c2, c4 and c5. Interestingly, 82 unique genes in 132 contrasts exhibited significant but opposite direction of expression change in PD between males and females (**Table S6**). For example, *SLC26A3* (solute carrier family 26 member 3), *RASGEF1B* (RasGEF Domain Family Member 1B), and *LINGO1* (Leucine Rich Repeat And Ig Domain Containing 1) were consistently down- regulated in male patients in every cluster, but up-regulated in female patients in several clusters (**Fig. S9B**). In contrast, *HBB* (hemoglobin subunit Beta), which was down-regulated in all but 5 clusters in female patients, was up-regulated in multiple clusters in the male patients (**Fig. S9B**). The gene ontology terms most enriched for those genes with opposite direction of expression change in PD between females and males were involved in cell morphogenesis, neuron development, and neuron differentiation (**Fig. S9C**), implicating an impact of sex on the gene expression for developmental and differentiation during PD progression.

### Cell type-specific gene expression enrichment and de-regulation of PD-associated genes in the SN

We next examined the expression of PD-linked genes and GWAS risk alleles in our dataset. We were able to detect the expression of 22 PD-linked genes ^3, 33^, of which half (11/22) are enriched in neuron clusters, such as GABAergic (c7_1, adjusted p=0.019, OR=17.9), glutamatergic (c7_2, adjusted p=1.5E-04, OR=23.0) and DA neurons (c7_3, adjusted p=9.1E-05, OR=25.5). *UCHL1 (PARK5)*, *SNCA (PARK1/4)*, *ATP13A2 (PARK9), VPS35 (PARK17)*, *SYNJ1 (PARK 20), CHCHD2 (PARK 22)*, and *TMEM230* showed strong expressions in both glutamatergic (c7_2) and DA (c7_3) subcluster neurons, while *PINK1 (PARK6)*, *EIF4G1* (*PARK18*) and *GBA* are particularly enriched in DA neuron subcluster (c7_3) (**Fig. 5A**, left). Unlike neuron-enriched PD genes, *LRRK2 (PARK8)* was highly expressed in microglia (c1), endothelial cells (c5) and OPCs (c4). *PRKN (PARK 2)* is enriched in astrocytes (c2), microglia (c1), oligodendrocytes (c3) and OPCs (c4). Furthermore, *VPS13C (PARK 23)* and *DNAJC13 (PARK 21)* were enriched in microglia (c1) (**Fig 5A,** left).

**Fig. 5.**
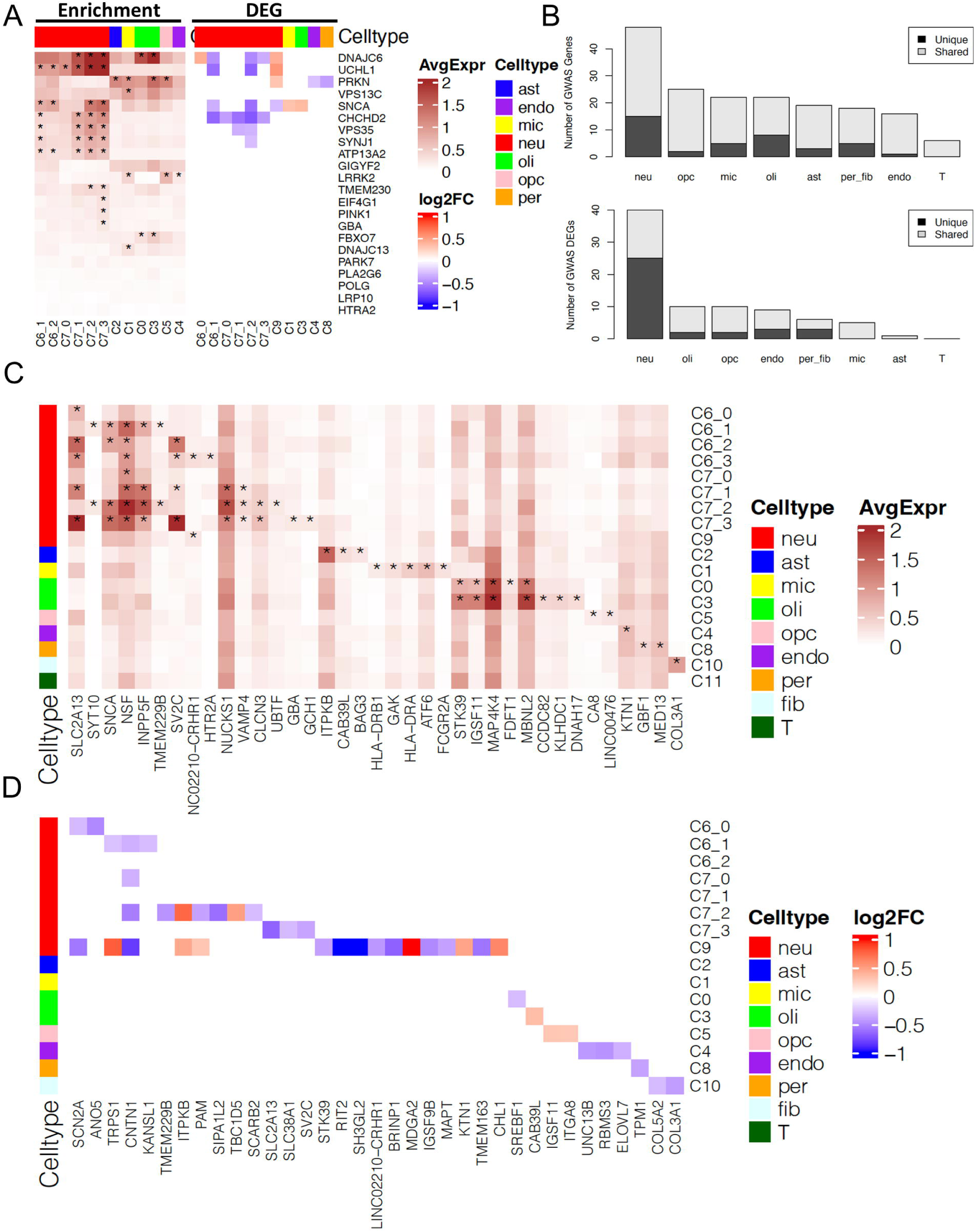
Cell-type-specific expression and DEG of PD-associated genes and risk loci. **A**, Heatmap for cell-type-specific expression (left) and DEG (right) of *PARK* family genes in PD. **B**, Number of GWAS loci related genes enriched (top) and differentially expressed (bottom) in each cell type. Dark grey indicated genes unique to each cell type and light grey indicated genes shared among cell clusters. **C**, Expression patterns of selected PD GWAS loci-related genes in different cell clusters. **D**, Heatmap for the log2 fold change (FC) of differential expression of selected PD GWAS loci genes in each cluster.

The differential regulation of PD genes is heterogeneous across cell types. Nearly 30% of the PD genes (6/22) were downregulated in neuronal clusters. For examples, we found that *DNAJC6*, *CHCHD2,* and *SNCA* were downregulated in DA neuron subclusters c7_3 or c9. In contrast, *DNAJC6*, *UCHL1* and *PRKN* were upregulated particularly in DA subcluster c9, whereas *SNCA* was upregulated in microglia and oligodendrocytes. In contrast, *PRKN* was downregulated in pericytes (c8) and endothelial cells (c5) in PD (**Fig 5A**, right).

We next extended our studies to the genes mapped to the known PD GWAS loci curated by GWAS catalog (www.ebi.ac.uk/gwas/, see Methods). Among 278 PD GWAS loci alleles, 90 genes showed cluster-specific expression as evidenced by significant enrichment in one DA neuron cluster (c9, adjusted p=0.037, OR=3.8), while another DA neuron cluster (c7_3) is enriched with genes linked to genetic forms of PD. 52 GWAS-associated genes were differentially expressed between PD and control (**Fig. 5B**). By categorizing these genes by cell type, we found that PD GWAS loci genes were preferentially expressed in neurons (**Fig. 5C**) and more frequently de- regulated in neurons than in other cell types (**Fig. 5D**). For example, *SV2C* is highly expressed in both c6_2 and c7_3 DA neuron clusters and downregulated in c7_3 in PD (**Fig. 5C and 5D**). *SV2C* encodes a synaptic vesicle glycoprotein 2C, which plays a role in the control of regulated secretion in neurons^34^. *KTN1*, encoding an integral membrane protein belonging to kinectin family, is highly expressed in endothelial cells (c4) (**Fig. 5C**) but upregulated in DA neurons (c9) (**Fig. 5D**). The above results demonstrate the heterogeneity of expression enrichment for PD-associated genes in different cell types of the SN, suggesting the complexity of pathogenic mechanisms of PD.

### DA neurons exhibit trajectories that correlate with PD progression

To further dissect how transcriptomes are altered in the SN during PD progression at the single-cell level, we performed pseudotime trajectory analysis of each cell cluster using a Monocle-based method^35^ (**Fig. 6** and **Fig. S9**). By trajectory analysis, we sorted cells along a learned trajectory, and their changes in the progress of trajectory were modeled with a dynamic function metric called pseudotime. Only DA neuron clusters (c7 and c9) showed mean trajectory pseudotime values in PD cells significantly higher than that in control cells at a cutoff of fold change ≥ 1.5 (t-test P value close to 0) (**Fig. S9A, Fig. 6A-B**). Trajectory inference assigned cells into multiple branches (or states) which formed a tree-like graph structure **(Fig. 6A** and **Fig. S9B**,). The paths traversing through connected branches may represent developmental/differentiation routes that cells could potentially go through (e.g., from healthy or progenitor into disease or terminally differentiated states). Therefore, we manually specified two longest potential routes starting from branches with lowest pseudotime to the branches with highest pseudotime for both c9 (**Fig. 6A**) and c7 (**Fig. S9D**), and further identified genes whose expression values were correlated with the pseudotime dynamics along the routes. We found a list of genes most strongly correlated with either or both routes, including *ZNF385D*, *FGF14*, *KCND2*, *CDH18*, *DHFR,* and *RASGEF1B*. *RIT2* and *SNCA* were also correlated with both routes (**Fig. 6C**). Further analysis revealed the similarity and discrepancy between neurons following different progression routes (**Fig. 6D**). For c9, similar gene ontology pathways, including neuronal system, neurexin and neuroligin signaling, and protein-protein interactions at synapses, were enriched in genes negatively correlated with both routes 1 and 2. In contrast, spinal cord injury, oligodendrocyte specification and differentiation, and neural crest differentiation were enriched in the genes positively correlated with route 2, while ERBB2 signaling and ferroptosis were enriched in the genes positively correlated with route 1 (**Fig. 6D**). For c7, neuronal pathways (e.g. neuronal development, axon, and synapse) were enriched in genes negatively correlated with both routes 1 and 2, especially route 2 (**Fig. S9E**). Pathways related to receptor inhibition and apoptosis were enriched in genes positively correlated with both routes 1 and 2, while myelin sheath and glial cell differentiation were only enriched in route 2 (**Fig. S9E**). The genes with the strongest correlation with trajectory routes include *DHFR*, *MAP1B*, *MTRNR2L8*, *IDS*, and *SNAP25 (***Fig. S9F**). These trajectory analysis results indicate heterogeneous cellular responses in the DA neurons during PD progression.

**Fig. 6.**
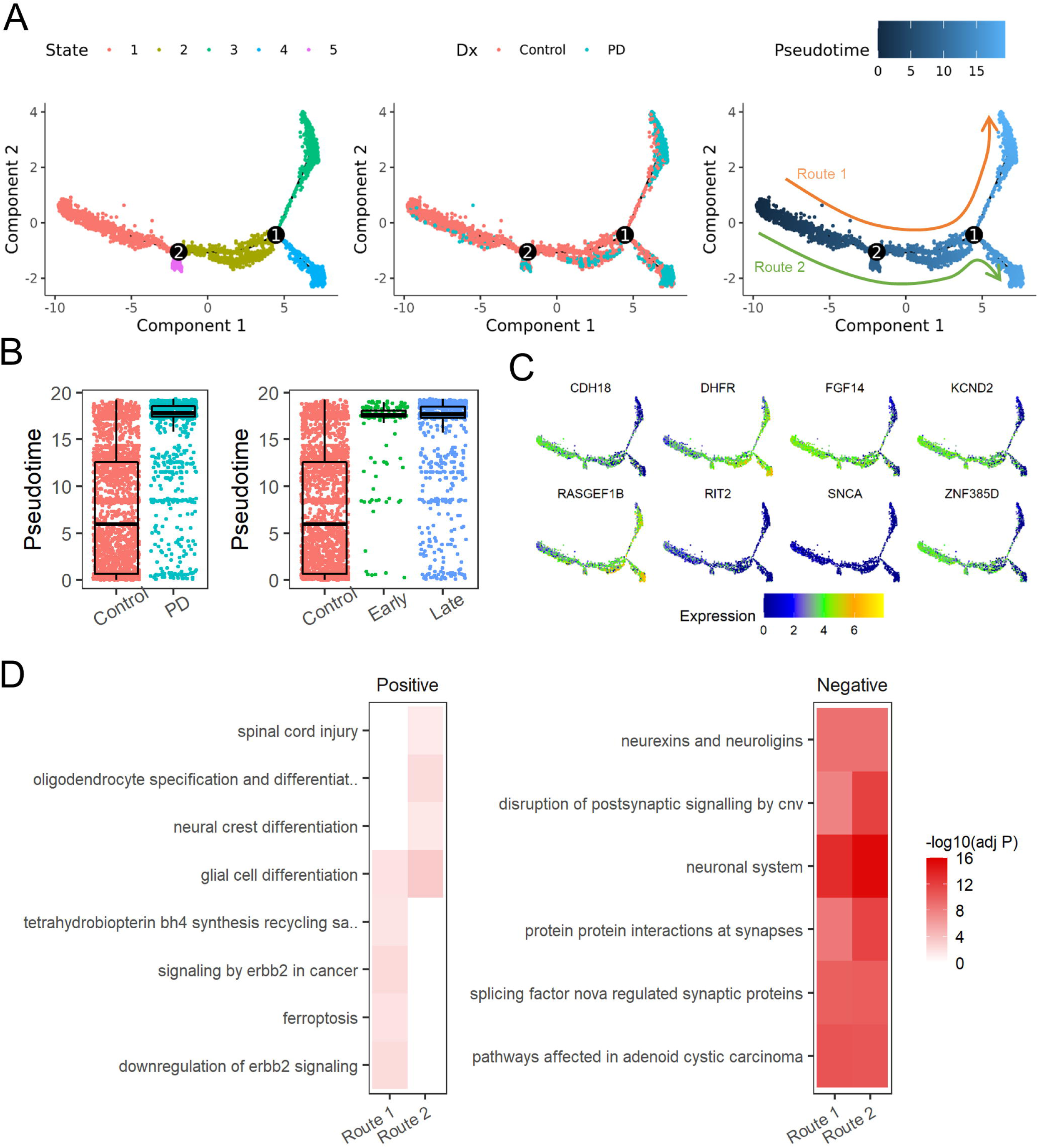
Trajectory inference of cluster c9 neurons. **A**, Cell distribution along trajectory in c9 by State (left), Disease status (middle) and Pseudotime (right). **B**, Comparison of pseudotime by PD and control (left) and by Braak stage(right). **C**, Selected genes whose expression significantly correlated with the inferred pseudotime in Cluster 13. **D**, Top pathways enriched in genes positively or negatively correlated with the pseudotime in two potential trajectory routes as indicated in (**A)**.

### Altered cell-cell communication networks in SN of PD

We next predicted altered cell-cell communications by using an R package CellChat, where cell communications are characterized by ligand-receptor (LR) interactions between the source and target cells^36^. Firstly, by comparing the differential number and strength of LR interactions among major cell types between control and PD, we observed a global decrease of cell communications for neuronal cells, but increased communications for microglia, pericytes, endothelial, and fibroblasts (**Fig. 7A**). Secondly, we assessed how individual cell clusters are affected by PD, according to the total outgoing and incoming signals. A number of cell clusters showed a loss of incoming and/or outgoing cell communications, including c0 (Oli), c2 (Ast), c3 (Oli), c5 (OPC), c6 (Neu), and c9 (Neu) (**Fig. 7B**).

**Fig. 7.**
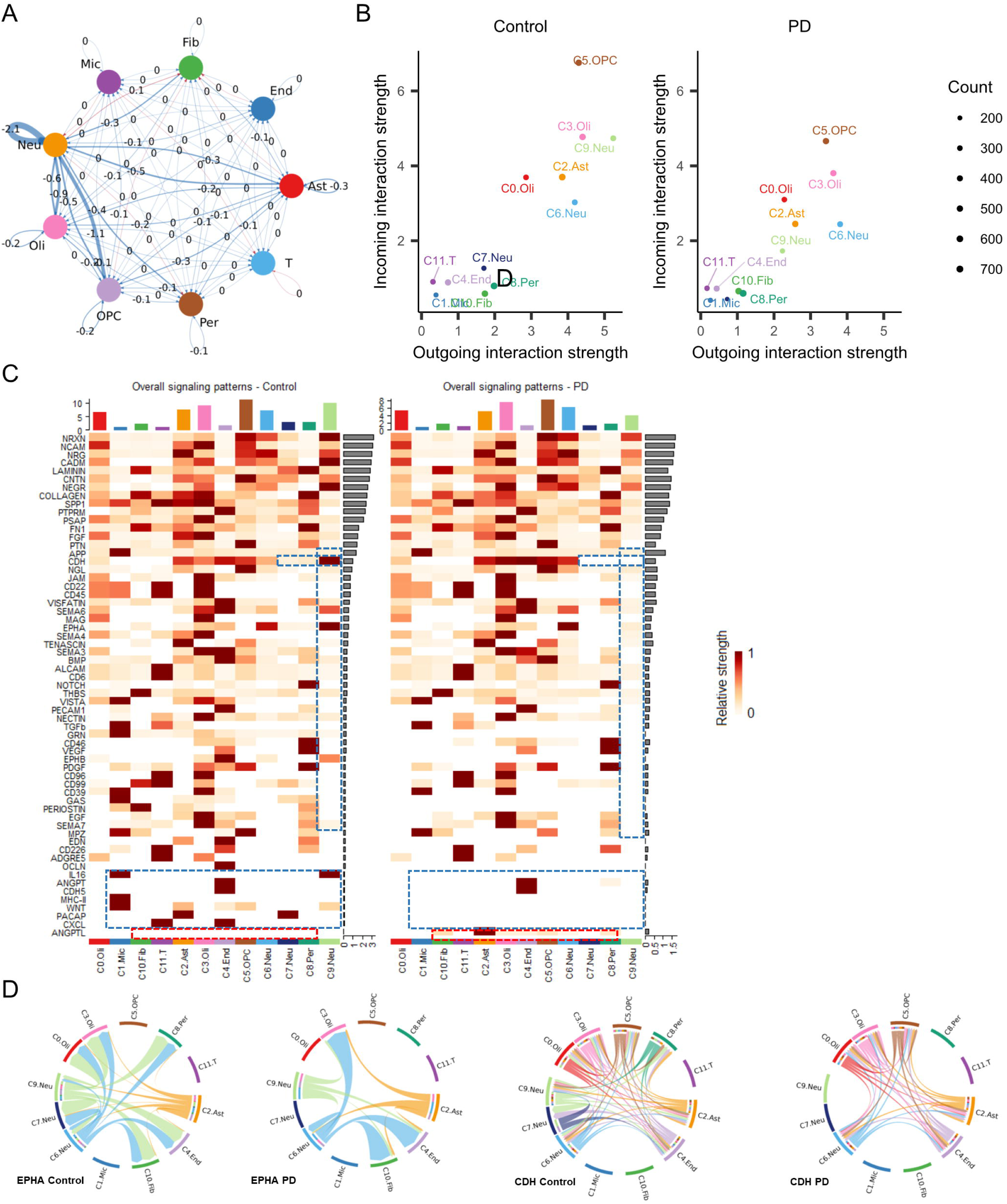
Altered cell communication networks in PD. **A**, Differential ligand-receptor interactions between PD and the controls. Blue lines and red lines indicated decreased and increased interactions, respectively. The line width was proportional to the difference. **B**, Distribution of cell cluster based on their relative incoming and outgoing signaling strength. **C**, Information flow changes of major signaling pathways between PD and control in each cell cluster. Dashed boxes highlighted gain (red) or loss (blue) of signaling in specific cell clusters. **D**, Chord diagram of EPHA and CDH signaling in control and PD.

By aggregating the LR pairs into major signaling pathways, we found that GRN, EPHB, GAS, PERIOSTIN, EDN, OCLN, IL16, MHC-II, WNT, PACAP, and CXCL pathways are inactivated in PD (**Fig. 7C**). Interestingly, the APP pathway was enhanced while the ANGPTL pathway was activated in PD. Multiple signaling pathways also showed cell cluster-specific regulation in PD. For example, two neuronal clusters c7 and c9, both containing subtypes of DA neurons, lost CDH signaling in PD (**Fig. 7C and 7D**). Disruption of CDH pathways was also seen in pericytes (c8). *CDH2*, which encodes N-cadherin, is the primary member of CDH. Because N-cadherin exerted a neuroprotective effect on DA neurons^37^, the loss of CDH interactions in DA neurons may disrupt the functions of DA neurons. Ephrins and Eph receptors are multifunctional in various biological conditions, including axon guidance and regeneration^38^. Aside from the loss of EPHB signaling in PD, we noticed that the EPHA input from neuronal clusters c6 and c9 into neuronal cluster c7, oligodendrocyte cluster c0, and pericytes cluster c8 are lost in PD (**Fig. 7D**). Taken together, our analysis suggests the alterations of multiple pathways in PD. Specifically, we found extensive disruptions and re-arrangements of CDH and EPHA/EPHB signaling pathways in the SN of PD.

## Discussion

Emerging sc/snRNA-seq approaches not only have become vital tools for dissecting heterogeneity and composition of cell types, but also provided significant insights into molecular mechanisms for complex human diseases such as Alzheimer’s disease (see our review)^39^ . Here we profiled 315,867 single nuclei, which is unprecedent, in the human SN from 32 postmortem brains including PD and controls of an average age at 81. We developed a comprehensive transcriptomic atlas of the human SN in a cell type specific manner. Importantly, our data reveals cell-type-specific molecular alterations and disruption of cell-cell communication networks in the SN of PD. Our studies highlight the cell heterogeneity and reveal molecular basis for vulnerability and potential resilience of human DA neurons in PD.

A significant finding from our study is the identification of three molecularly distinct subtypes of DA neurons in human aged SN. The classification of human DA neurons in our study shows clear distinction from that of the previous mouse midbrain DA neurons, which are largely based on the TFs determining the formation and differentiation of DA neurons during mouse brain development^25^. While the three human DA neuron subtypes degenerate in PD, the composition of the rest cell types remains relatively unaltered in the SN. By integrating snRNAseq data from two independent reports and IHC validation using separate cohorts in our study, we identified a *RIT2*- enriched DA neuron subtype (*RIT2^high^*) with reduced *TH* levels even in non-PD SN. In contrast, the other two subtypes of DA neurons express high levels of DA markers, such as *TH*, *SLC18A2,* and *SLC6A3*, and they are distinguished by differential expression multiple marker genes including *GALNTL6* and *DCC*. The finding of the *RIT2^high^* DA neuron subtype is unexpected. *RIT2^high^* neurons display significantly different gene expression pattern from two other DA subtypes (*TH^high^*). Yet *TH^high^* and *RIT2^high^* subtypes both appear to be associated with NM and degenerate in PD. A previous study found differential rate of degeneration between NM+ and TH+ neurons in the SNpc of PD, suggesting the heterogeneity of NM+ neurons - not all NM+ neurons express TH^17^. The identification of RIT2^High^/NM+ populations can explain the observation of NM+TH- neurons ^17^ and supports the idea that *RIT2*^High^/NM+ is originated from DA neuron, as the formation of NM requires L-DOPA, a precursor of DA synthesized by TH. It is conceivable that the *RIT2*^high^ DA neurons reduce or lose typical DA markers such as *TH*, *SLC18A2* and *SLC6A3* production but retain NM during aging process. The lack of significant *TH, SLC18A2* and *SLC6A3* expression suggests that *RIT2*^high^ neurons are functionally inert as DA neurons (“ghost”) and/or may develop distinct functions in the SN of aged individuals.

While the emergence and biological function of *RIT2*^high^ neurons remains to be elucidated in the future, *RIT2* was previously identified as a PD susceptible gene^23^. RIT2 protein belongs to the RAS superfamily of small GTPases, which interacts and regulate DAT levels in a sex-dependent manner in mice ^40^. Furthermore, knockdown of *rit2* in DA neurons impacts differential DA-dependent behavior in male vs. female mice^40^, demonstrating a role for RIT2 in regulating specific DA neuron functions. We found *RIT2* expression was down-regulated in the subtype of DA neurons (c9) in PD with advanced Braak stages (0.8-fold, adjusted P value=0.047), while a previous report also showed reduced expression of *RIT2* in the SN of PD brain^41^. Therefore, the above evidence raises a possibility that *RIT2*^high^ DA neurons are molecularly and functionally distinct from *TH^high^* DA neurons in the SN; loss of *RIT2*^high^ neurons may contribute to certain symptoms of PD that should be investigated. We propose that *RIT2* could serve as a potential marker in age-related pathological diagnosis of PD due to the loss of *RIT2*^high^ neurons additional to *TH^high^* neurons in the SN of PD.

The PD samples used in our snRNAseq study are from advanced stage of PD (∼80), where the majority DA neuron population are lost. Therefore, our data is unlikely to provide an insight into molecular events at the initiation and during progression of neurodegeneration. However, we theorize that study of the remaining DA neurons at advanced stage helps reveal a mechanism whereby they are adapted and acquire resilience. In fact, *Kordower et al* reported a rapid decline of DA neuron number in the SNpc of PD at early years of symptom, while a subpopulation of DA neurons remained unchanged in number through at least a decade at advanced stage of the disease ^17^.

At present it is entirely unknown what could render the remaining DA neuron resistant to the death at late PD. By comparing transcriptomic profiles of the DA neurons between PD and control cohorts, our study provides an opportunity to decipher potential mechanism for their resilience. For instance, apart from upregulation of ribosomal genes, protein translation-related pathways, heat-shock related genes, and metallothionein family genes common among many cell types in PD, DA neuron subtypes share the up-regulation of several genes associated with DA neuron protections. PLCL1 is a phospholipase C-related inactive protein type 1, which is upregulated in all three subtypes of DA neuron of PD. PLCL1 is shown to regulate GABA receptor trafficking and PLCL1 knockout mice display motor function deficits ^42, 43^. *CST3, TMEFF2, and CRYAB* are increased in at least two DA neuron subtypes of PD. *CST3* is an inhibitor of cathepsin B and was shown increased in the DA neurons of PD through laser capture analysis ^44^, and it plays a role in protecting DA neurons ^45^. *TMEFF2* is a transmembrane protein with EGF like and two follistatin like domains 2 and considered a survival factor for hippocampal and dopaminergic mesencephalic neurons ^46^. In addition, altered *CRYAB* levels may impact autophagic activity and the clearance of α-synuclein according to previous studies ^29, 30^. The above notions warrant future experimental validations.

Our study gains an insight of the cell-type specific gene expression for the PD-linked genes and GWAS risk alleles. Our results demonstrate the cell heterogeneity of expression enrichment for PD-associated genes in the SN. The DEG analysis suggests common and distinct cellular pathways that are affected in various cell types of the SN of PD. Our result reveals that the majority of DEGs in PD are associated with neurons. It is worth noting that *LRRK2* is produced primarily in microglia and OPC, but little change was observed in any cell-type in PD. Furthermore, *SNCA* expression is reduced in subpopulations of DA and glutamatergic neurons however enhanced in microglia as well as oligodendrocyte. The transcriptional elevation of *SNCA* has not been previously reported in glia at the SN of PD, and the significance of this observation should be investigated in the future. The above observations highlight the diversity of the molecular mechanisms underlying DA neuron degeneration.

Many studies have indicated activation of microglia or astrocytes in PD. Several groups reported the appearance of amoeboid microglia producing MHC class II, ICAM-1, and LFA-1, the markers for activated microglia, and reactive astrocyte expressing ICAM-1 in the SNpc and the putamen of PD brain^47–51^. Previous studies also detected increased binding of a radiotracer, 11C- (R)-PK11195, in PD brains compared to controls. This tracer is known to bind to 18-kDa translocator protein (TSPO) expressed mainly by microglia^52^. However, other groups failed to observe activated microglia or astrocytes in human PD brains^53, 54^. In our study, we found no clear evidence supporting the activation of inflammation-related molecules or disease-associated microglia signature in glial clusters, although we detected the up-regulation of multiple genes, such as *AKT-PIP3, FOXO1/3* and *ERBIN,* and *NGR3,* which are known to regulate macrophage/microglia activation, migration, proliferation and inflammation^55–59^. The discrepancy of the results could be due to many factors such as sample source and detection of procedures; it remains possible that glia become less active at advanced stage of PD as examined in our study. Furthermore, accumulated evidence also suggests the increase of senescent/dystrophic microglia in human aged brains^60, 61^.

We note that our study has several caveats. Despite the largest PD samples ever analyzed by snRNAseq, the sample size of our study is still relatively small, considering the variations among postmortem samples such as PMI and pathological differences. The observation of altered gene expression in PD could be biased and should be rigorously validated with large sample sizes from independent cohorts. Moreover, the emergence and physiology of *RIT2*-enriched neurons in the human SN has yet to be elucidated.

In summary, our study has established a transcriptomic atlas of the human SN at single-cell resolution and delineated the landscape of molecular and cellular alterations in PD. Our study not only provides a valuable resource for dissecting molecular and cellular compositions and structures of the human SN, but also presents an unprecedented opportunity to understand in-depth pathogenic mechanisms, identify key therapeutic targets and develop clinical biomarkers for PD.

## Materials and Methods

### Postmortem brain sample collection

The postmortem brain samples were requested from NIH Neurobiobank (www.neurobiobank.nih.gov) and fulfilled by Brain Endowment Bank at Miller School of Medicine, University of Miami. The samples were pre-tested for known genetic mutations linked to familial PD, including *SNCA*, *LRRK2* and *GBA*. The samples did not harbor any of the abovementioned mutations.

The information of the sample demographics was summarized in **Table 1** and **Table S1**. Specifically, frozen punches of substantia nigra were obtained and then pulverized in a liquid-nitrogen-chilled mortar and aliquoted. Approximately 50mg tissues were used for snRNA-seq.

### Nuclei isolation and sequencing

Single nucleus gene expression sequencing was performed on the samples using the Chromium platform (10x Genomics, Pleasanton, CA) with the Next GEM Single cell 3’GEX Reagent kit, and an input of ∼10,000 nuclei from a debris-free suspension. Briefly, nuclei were isolated from frozen tissue, as per 10x Genomics’ recommendations, using chilled, 0.1% Nonident P40 lysis buffer with gentle homogenization and washed. Gel-Bead in Emulsions (GEMs) were generated on the sample chip in the Chromium controller. Barcoded cDNA was extracted from the GEMs by Post-GEM RT-cleanup and amplified for 12 cycles. Amplified cDNA was fragmented and subjected to end-repair, poly A-tailing, adapter ligation, and 10X-specific sample indexing following the manufacturer’s protocol. Libraries were quantified using Bioanalyzer (Agilent) and QuBit (Thermofisher) analysis. Libraries were sequenced using a 2x100PE configuration on a NovaSeq instrument (Illumina, San Diego, CA), targeting a depth of 50,000-100,000 reads per nucleus.

Sequencing data were aligned and quantified using the Cell Ranger Single-Cell Software Suite (version 3.1.0, 10x Genomics) against the provided GRCh38 reference genome using default parameters, including introns. Before sequencing all samples, 3 samples were randomly selected for sequencing in a pilot run. Then all 31 samples were processed and sequenced in one batch. For the 3 samples with replicated libraries, we observed similar data quality between the pilot and final sequencing run. Therefore, we combined libraries from both the pilot and final sequencing, resulting in a total number of 457,453 nuclei (on average 13,455 nuclei per library) before quality control (QC).

### snRNA-seq data preprocessing and pre-clustering analysis

Starting from Cell Ranger derived unique molecular identifier (UMI) count matrices from all sequencing libraries, we performed QC by removing low-quality nuclei with either too few genes (< 200) or an excessive number (> 2500) of genes detected, retaining 355,157 nuclei after filtering. Then we removed insufficiently detected genes by keeping 30,038 genes expressed in more than one nucleus. Mitochondrial reads were discarded to avoid biases introduced during the nuclei isolation since they are not expressed inside nucleus^62–64^. After QC, we obtained on average 11,099 nuclei per individual, and 817 unique genes per nuclei per individual. We performed a pre-clustering analysis using a well-established scRNA-seq data integration workflow based on R packages Harmony^19^ and Seurat (v3)^18^. Briefly, the UMIs data was first normalized by sequencing depth and log-transformed using the LogNormalize method implemented in Seurat. 2,000 most variable gene features were identified, scaled and centered after regression out covariates sex, age, and postmortem interval (PMI). Next, dimensional reduction was performed using principal component analysis (PCA) based on the 2,000 most variable genes. The top 30 principal components (collectively explaining more than 90% of the variance) as determined by an elbow approach were selected for integration of snRNA-seq data across all sequencing libraries with Harmony^19^. Top 20 embeddings in the Harmony space were used for calculating 2D dimensional reductions by t-distributed Stochastic Neighbor Embedding (t-SNE)^65^ and Uniform Manifold Approximation and Projection for Dimension Reduction (UMAP)^66^. The same top 20 Harmony embeddings were also used to compute the nearest neighbor graph and the subsequent cell pre-clusters with the Louvain algorithm implemented in Seurat^18^. This initial pre-clustering analysis resulted in 14 pre-clusters at resolution 0.2. Two smallest pre-clusters c12 and c13, dominated by nuclei from one or two donors, overlapped with pre-cluster c0 on the UMAP space (Fig. S1A-B).

### Cluster stability analysis

To assess the stability and robustness of the pre-clusters, we performed repeated subsampling analysis by making use of software tool scclusteval^67^. In each subsample, we sampled without replacement a subset of 80% of the nuclei in the full QCed dataset, and then repeated the data normalization, scaling, PCA, Harmony data integration, and clustering procedure on this subset of data as above described. We repeated subsampling 100 times. For each subsample, we compared its clusters with those pre-clusters from the full data by Jaccard index analysis and returned a maximum Jaccard index coefficient for each of the original pre-clusters. We found that pre-cluster c12 had almost close to 0 Jaccard index coefficients in all subsamples, indicating that it was an unstable cluster dissolved in the subsamples. Pre-cluster c13 was also dissolved in 13 of the subsamples, suggesting it was a potentially unstable cluster. For the remaining pre-clusters, they all showed Jaccard index coefficients larger than 0.50, except for c9 where one subsample had a Jaccard index coefficient of less than 0.25. Next, for each cell *i* in pre-cluster c12 or c13, we assessed the co-clustering probability between *i* and all the pre-clusters as the mean fraction of cells in the pre-clusters that clustered together with *i* in the repeated subsamples by using equation:

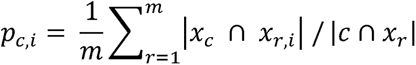

where *m* denotes the number of subsamples that included cell *i*, *c* denotes a pre-cluster, *x_c_* denotes the set of cells in pre-cluster *c*, *x_r_* denotes the set of cells in subsample *r*, and *x_r,i_*, denotes the set of cell subcluster that contains cell *i* in *x_r_* The distribution of *p_c,i_*, stratified by *c* is shown in Fig. S1D. Cells in pre-cluster C12 tended to co-cluster with cells in pre-cluster C0, followed by cells in C3 and C13, while cells in pre-cluster C13 tended to co-cluster with cells in pre-cluster C12, followed by cells in C0. Together with the spatial distribution of the cells in the UMAP space, we decided to merge the two unstable pre-clusters C12 and C13 into their adjacent bigger neighbor pre-cluster C0, leading to 12 clusters (C0-11) for further analysis.

### Doublet prediction analysis

After finalizing the cell clusters, we predicted doublets by making use of the scDblFinder package^68, 69^. scDblFinder first simulates doublets from the provided cell clusters and then computes a doublet prediction score for each cell by combining the fraction of simulated doublets in its neighborhood with another score based on co-expression of mutually exclusive gene pairs^70^. The doublet prediction scores is iteratively refined and a classification model is trained to best characterize the putative doublets by integrating a number of discriminating metrics^68, 69^. Fig. S1E shows the final doublet prediction score and classification. 39,290 predicted doublets were removed, resulting in 315,867 singlets the final clusters for all downstream analyses.

### Cluster cell type annotation

For each major or sub- cluster, we first interrogated the expression patterns of known gene markers to annotate clusters into major cell-types: neurons (*RBFOX3*, *GAD1*, *NRGN*), astrocytes (*AQP4*, *GFAP*), oligodendrocytes (*MOG*), microglia (*C3*, *CSF1R*, *CD74*, *TYROBP*), oligodendrocyte progenitor cells (*VCAN*), endothelial cells (*FLT1*), and pericytes (*PDGFRB*). Next, we calculated *de novo* cluster signatures by comparing the cells in this cluster against the cells of the rest clusters using Wilcox rank sum test in Seurat. We defined cluster up-regulated genes as those up-regulated by at least 1.2-fold and with Bonferroni adjusted P value less than 0.05, from which we further defined cluster-enriched *de novo* signatures (i.e. marker genes) as those up-regulated by at least 2- fold. To assist the annotation of cell type of each cluster, we overlapped the *de novo* cluster signatures with a large-scale collection of cell type markers curated from over 1,054 single-cell experiments^20^, with P value significance of the overlaps computed by a hypergeometric test. Since cell type marker expression may change in PD cells, only the control cells were used for investigating the marker gene expression pattern as well as calculating the cluster signatures.

### Sub-clustering analysis

To perform sub-clustering analysis for a given cluster, we first extracted the normalized and covariates adjusted data for the cluster. As in the pre-clustering analysis, we computed dimensional reduction using PCA. Since the number of cells contributed from each individual donor ranged from 18 to 2,170 (cluster 6), and 19 to 2,069 (cluster 7), we did not conduct Harmony analysis to avoid biased data integration due to small and uneven cell numbers. The top 10 principal components as determined by an elbow approach were selected to compute UMAP, the nearest neighbor graph and the subsequent cell sub-clusters with the Louvain algorithm. Sub-cluster marker signatures were defined by comparing each sub-cluster with all other cells, including other sub- clusters and major clusters, using Wilcox rank sum test in Seurat as above.

### Cell cluster proportion change

To test if there was a significant change in the proportion of a cell cluster between PD and controls, we first calculated the odds ratio (OR) of proportion difference using formula: 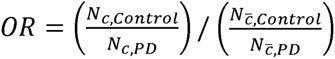, where *N_c,PD_* (*N_c,Control_*) denotes the total number of cells in cluster *c* in all PD (Control) samples, while 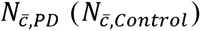 denotes the total number of cells in the remaining clusters in all PD (Control) samples. To compute the P value significance, we computed the *odds* of cells assigned to cluster *c* in individual *i* as 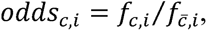 where *f_c,i_*, denotes the fraction of cluster *c* cells in the i-th individual and 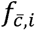, denotes the fraction of cells in all the other clusters in the *i*-th individual. Then a one-tailed Wilcox rank sum test was conducted to compare the difference in *odds* between PD and controls.

### Replication of cell clusters in two independent non-PD SN samples

To replicate the present cell clusters, we compared our data with two published snRNAseq datasets from non-PD SN samples. We first reprocessed the snRNA-seq data reported by Agarwal and colleagues from non-PD SN samples^71^. By using the pipeline described in the pre-cluster analysis section, we identified clusters of major brain cell types from the Agarwal data, including astrocytes, endothelial, microglia, neurons, oligodendrocytes, and oligodendrocyte progenitor cells (**Fig. S4A** and **S4C**). We further performed a sub-clustering analysis on the neuronal cluster c6 and identified 3 sub-clusters, c6_0 (*RIT2* and *RBFOX3* enriched), c6_1 (*TH* enriched), and c6_2 (*GAD1* and *GAD2* enriched) (**Fig. S4B** and **S4D**). To compare the cluster similarity between our data and the Agarwal data, we assessed the significance of intersection of cluster signatures between ours and the Agarwal dataset using hypergeometric test (**Fig. S4E**).

Similarly, we reprocessed another snRNA-seq data from non-PD SN samples reported by Welch et al.^14^ and identified clusters of major brain cell types (**Fig. S5A** and **S5C**). We performed a sub- clustering analysis on the neuronal cluster c4 and identified 7 sub-clusters, among which c4_3 is enriched for *TH*/*SLC6A3*/*SLC18A2* and c4_5 is uniquely enriched for both *RIT2* and *RBFOX3* (**Fig. S5B** and **S5D**). Then we compared the cluster signature similarity between our data and the Welch data using hypergeometric test (**Fig. S5E**).

### Cluster-specific differential gene expression and functional enrichment analysis

DEGs between PD and control and DEGs between different Braak stages and control in each cluster/sub-cluster were identified using MAST^72, 73^ implemented in Seurat, with correction for PMI, age, and sex. Sex-specific DEG analysis was performed using the same pipeline for each sex group separately, with PMI and age corrected. DEGs were identified at the cutoff of Bonferroni corrected P value ≤ 0.05 and fold change ≥ 20%. For Braak stage dependent differential expression analysis, we separated the cells into three groups by the Braak stage of the donors: Control (Braak = 0), Early Stage (Braak 1-3), and Late Stage (Braak 4-6). Genes significantly increased/decreased in the Early-vs-Control AND Late-vs-Control contrasts were considered as positive/negative early and sustained responders. Genes significantly increased in the Early-vs-Control contrast but decreased in the Late-vs-Early contrast and vice versa were defined as positive and negative “U-shaped” responders, respectively. Genes significantly upregulated/downregulated only in the Late-vs-Control or also in the Late-vs-Early contrast were late responders. Functional enrichment of DEGs with MSigDB gene annotation collections was examined by Fisher’s Exact Test (FET) with Benjamini-Hochberg (BH) correction. Results with BH adjusted p value < 0.05 were considered statistically significant.

### Comparisons between cluster-specific with bulk-tissue-based DEGs

Bulk-tissue-based DEGs were defined by a meta-analysis described in our previous work^31^. The intersection between the present cell cluster-specific DEGs and the bulk-tissue-based DEGs was examined by FET followed by BH correction using the common genes identified in both bulk and snRNA-seq as the background. Results with BH adjusted p value < 0.05 were considered statistically significant.

### PD-associated gene expression and regulation patterns in the snRNA-seq

PD causal genes were defined as genes whose mutations were directly linked to familial PD as reviewed in ^3, 33^. PD GWAS genes were downloaded from GWAS catalog (www.ebi.ac.uk/gwas/) and defined based on the mapped genes closest to the risk loci. We first examined whether PD-associated genes were preferentially expressed in certain cell types by overlapping them with top 10% up-regulated genes ranked by fold change in each cell cluster compared to the rest using control cells only. The overlap was tested by FET using GeneOverlap package in R and the results with Bonferroni adjusted p value < 0.05 were considered statistically significant. We then examined whether PD-associated genes were differentially regulated in PD by overlapping them with cluster-specific PD-vs-Control DEGs. Total number of PD-associated genes that showed cell-type-specific enrichment and differential regulation were summarized and they were further separated into single cell-type enriched or shared across multiple cell types.

### Trajectory inference

The trajectory inference was performed for each cell cluster using a Monocle^74^ based script as previously described^35^. In this analysis, normalized gene expression data were extracted for each cluster and only genes with expression in more than 20% of the cells in the cluster were selected as features for trajectory inference. After a trajectory was learnt, cells were assigned into different branches (or states). Then we computed the cell changes along the trajectory with a dynamic function metric termed pseudotime. To define the beginning of the trajectory for pseudotime estimation, we selected the state containing most of the control cells as the root state. After pseudotime calculation, we compared the pseudotime values between control and PD cells and identified the clusters with significant pseudotime difference by at least 1.5-fold (adjusted P value < 0.05 by t-test) for further gene-pseudotime path correlation analysis by Spearman’s correlation.

Functional enrichment of pseudotime path correlated genes with MSigDB was examined by FET with BH correction. Results with BH adjusted p value < 0.05 were considered statistically significant.

### Cell communication analysis

Cell communication analysis and visualization were performed using the default setting in CellChat package^36^. All ligand-receptor pairs in cell-cell contact, extracellular matrix (ECM) receptors and secreted signaling were included. Specifically, we focused on the gain or loss of interactions in general cell types, the shift of the outgoing and incoming interactions, and the top signaling pathways altered among cell clusters between PD and control.

### Immunostaining for the human brain

FFPE blocks were sectioned into 5μm slices, deparaffinized with Xylene for 5minutes three times, and rehydrated with gradient EtOH (100%, 100%, 75%, 50%, and water). Antigen retrieval was performed in SignalStain® Citrate Unmasking Solution (14746, Cell Signaling Technology, MA) with a pressure cooker (Presto Electronic, WI) for 15 min. Endogenous peroxidase activity is blocked by 3% hydrogen peroxidase for 10 min. For RIT2 staining, RIT2 antibody (CF501757, OTI3F4 clone, ThermoFisher), ImmPRESS® HRP Horse Anti-Mouse IgG PLUS Polymer Kit (Peroxidase, MP7452, Vector laboratory, CA), and ImmPACT® VIP (peroxidase substrate, SK- 4605) were applied. For Tyrosine hydroxylase (TH) staining, TH antibody (AB152, Millipore), ImmPRESS®-AP Horse Anti-Rabbit IgG Polymer Kit (alkaline phosphatase, MP5401, Vector laboratory, CA), and Vector® Blue (alkaline phosphatase substrate) were applied. VGLUT2 were stained with VGLUT2 antibody (ZMS1026, Sigma), ImmPRESS®-AP Horse Anti-Mouse IgG Polymer Kit (alkaline phosphatase, MP5402, Vector laboratory, CA), and Vector® Blue (alkaline phosphatase substrate) were used according to the manufacturer’s instructions. After staining, brain slices were dehydrated by 100% EtOH followed by Xylene and mounted with VectaMount® Permanent Mounting Medium (H-5000-60, Vector laboratory, CA). Images were taken by Axiophot 2 (Zeiss, CA).

### Statistical Analysis

The data analyses were performed using R/4.0.3 and GraphPad Prism 9 (GraphPad Software, CA, USA). For demographic information, the results were represented as Mean±standard deviation (SD) for continuous and N(%) for discrete variables, respectively. For immunohistochemistry, the results were reported as Mean±standard error of the mean (SEM). The statistical significance of differences between two groups was determined using the unpaired two-tailed Student’s t test. p value<0.05 was considered statistically significant.

## Acknowledgments

ZY grant support: P20NS123220; R21NS109895; Dr. George Heaton for comments (Yue lab). KF grant support: F32AG056098

## Author Contributions

ZY and BZ conceived the project. QW initiated the project and performed data analysis. MW performed data QC and clustering analysis. IC performed validation experiments. KF and JFC contributed to validation experiments. LH prepared brain samples for sequencing. KB and RS provided support in snRNAseq. DD and XS provided brain samples and related information. ZY, QW, MW, IC and BZ wrote the manuscript. LH, KB, RS, KF, JFC, DD and XS provided comments and edits. All authors approved the manuscript.

## Conflict of Interest

The authors declared no conflict of interest.

## Supplementary information index

### Supplementary Figures

**Fig. S1. PD snRNA-seq analysis. A**, UMAP for the pre-clusters split by control and PD. **B**, Stacked barplot for the contribution of cells from individual donors in each pre-cluster. **C**, Pre-cluster preservation in the subsampling analysis as assessed by Jaccard index. **D**, Co-cluster probability for cells in the pre-cluster c12 and c13. **E**, Doublet prediction score and classification by scDblFinder.

**Fig. S2 Cell cluster *de novo* markers and verification of VGLUT2 expression in controls. A**. Heatmap for the top 5 signature genes identified in the control cells of each cluster. **B.** VGLUT2 (SLC17A6) staining in unaffected control brain. Paraffin-embedded sections containing substantia nigra pars compacta (SNpc) were stained with antibodies against VGLUT2 (blue). Scale bar, 100µm in the left panel; 50µm in the right panel.

**Fig. S3. Cluster cell type annotation using cell type marker genes from a public database.** Heatmap for the P value significance of enrichment for known cell type marker in the present cluster signatures.

**Fig. S4. Replication of cell clusters in an independent non-PD snRNA-seq data reported by Agarwal et al. A**, UMAP for the cell clusters in the Agarwal et al. data. **B**, UMAP for the sub-clusters of the neuronal cluster C6 of Agarwal et al. data. **C**, Violin plot for the expression pattern of known cell type markers in **A**. **D**, Violin plot for the expression pattern of known brain cell type markers in the neuronal sub-clusters in **B**. **E**, Heatmap for the P value significance of the intersection of the cluster signatures between our data and the Agarwal et al. data.

**Fig. S5. Replication of cell clusters in an independent non-PD snRNA-seq data reported by Welch et al. A**, UMAP for the cell clusters in the Welch et al. data. **B**, UMAP for the sub-clusters of the neuronal cluster C4 in the Welch et al. data. **C**, Violin plot for the expression pattern of known cell type markers in **A**. **D**, Violin plot for the expression pattern of known brain cell type markers in the neuronal sub-clusters in **B**. **E**, Heatmap for the P value significance of the intersection of the cluster signatures between our data and the Welch et al. data.

**Fig. S6. Comparison of the 3 DA neuron clusters. A,** Violin plots for the expression of top marker genes of DA neuron clusters. Color denotes the source cluster of the markers (red, c6_2; green, c7_3; blue, c9). **B**-**C**, Intersection of PD-vs-Control DEGs among 3 DA neuron clusters. **B**, Barplot for the significance of the number of overlapping PD DEGs among 3 DA neuron clusters C6_2, C7_3, and C9. The matrix of solid and empty circles at the bottom illustrates the “presence” (solid green) or “absence” (empty) of the DEG sets in each intersection. The numbers to the right of the matrix are DEG set sizes. The colored bars on the top of the matrix represent the intersection sizes with the color intensity showing the P value significance. The genes in the intersections of size ≤ 13 are shown above the bar while the genes in the intersections of size > 13 are shown in **C**. The present multi-set intersection analysis as well as the visualization are performed by R package SuperExactTest.

**Fig. S7. Expression pattern of known DA markers and TFs. A-B**, Violin plots for the expression of known DA markers and TFs in control (**A**) and PD (**B**) cells. **C**, Percentage (%) of cells expressing each of the known DA markers and TFs in clusters c6_2, c7_3, and c9.

**Fig. S8. Braak-stage-dependent DEG analysis. A**, Top pathways associated with early and sustained responding genes. **B**, Top pathways associated with “U-shaped” responding genes. **C**, Top pathways associated with late responding genes

**Fig. S9. Sex-dependent differential expression analysis. A**, Number of DEGs in each cell cluster between PD and control in males and females. **B**, Heatmap for the the expression pattern of top 50 DEGs with the largest fold change (FC) in females or males. **C**, Barplot for the top 30 gene ontology pathways enriched in the genes with opposite direction of expression change in PD between females and males.

**Fig. S10. Trajectory analysis. A**, Boxplot for the distribution of estimated trajectory pseudotime stratified by disease status in each cluster. **B-F**, Summary of trajectory analysis for cluster C7. **B-D**, Dimensional reduction plots showing the cell states computed from the trajectory analysis (**B**), the disease status of the cell donors (**C**), and estimated trajectory pseudotime (**D**), with two potential cell progression routes highlighted by arrowed curves in **D**. **E**, Heatmap for the top gene ontology and pathways enriched in the genes positively or negatively correlated with the trajectory routes 1 and 2. **F**, Dimensional reduction plot showing the expression pattern of 10 genes mostly correlated with the trajectory routes.

### Supplementary Tables

Table S1 Sample demographic information.

Table S2 Cluster marker genes.

Table S3. Cluster-specific DEGs between PD and control.

Table S4. Braak stage dependent DEGs.

Table S5. Sex-dependent PD-vs-Control DEGs.

Table S6. DEGs with opposite expression changes between males and females.

## Notes

### Competing Interest Statement

The authors have declared no competing interest.

