## Supplementary figures and images for "Single-cell transcriptomic atlas of the human substantia nigra in Parkinson’s disease"

### FIG S1

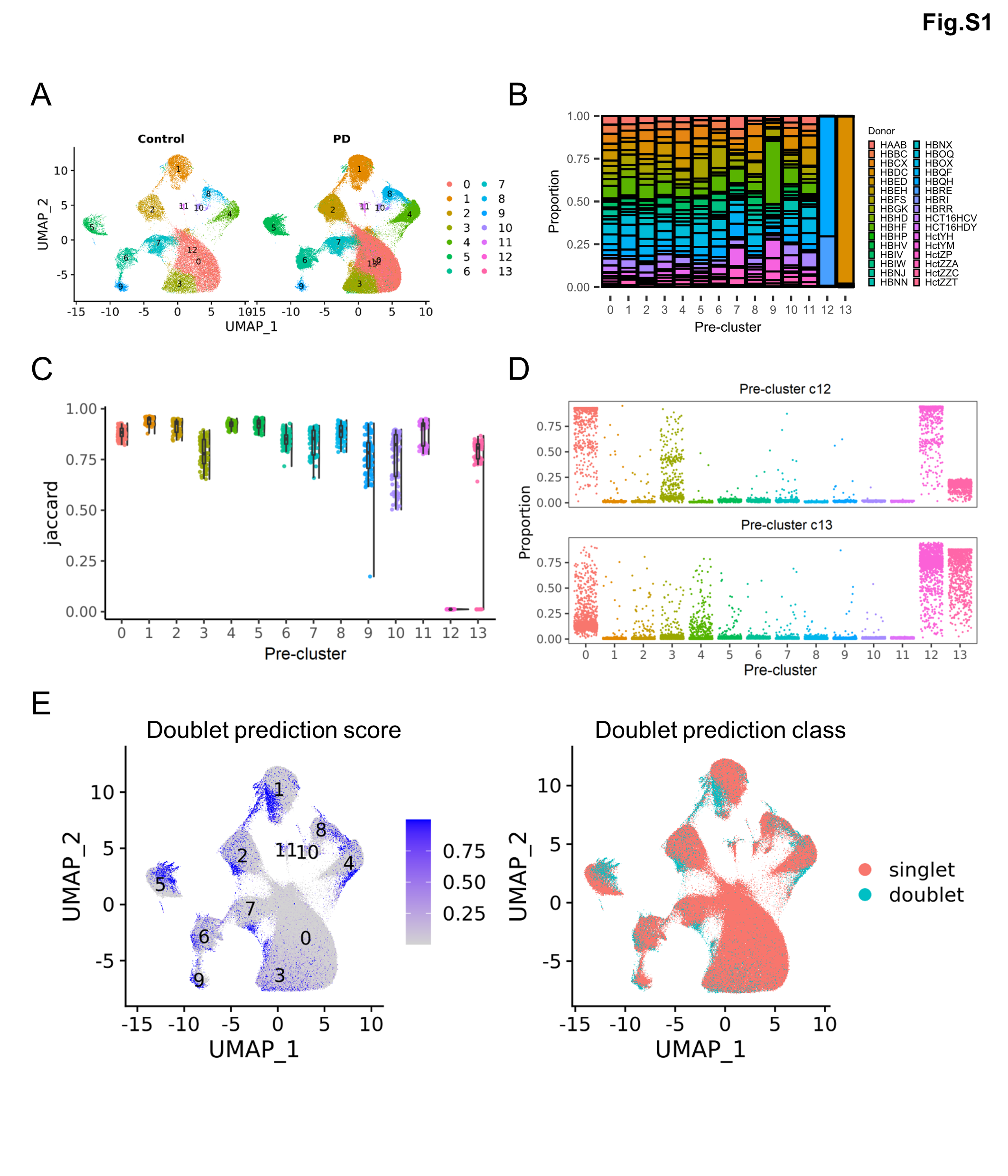

### FIG S2

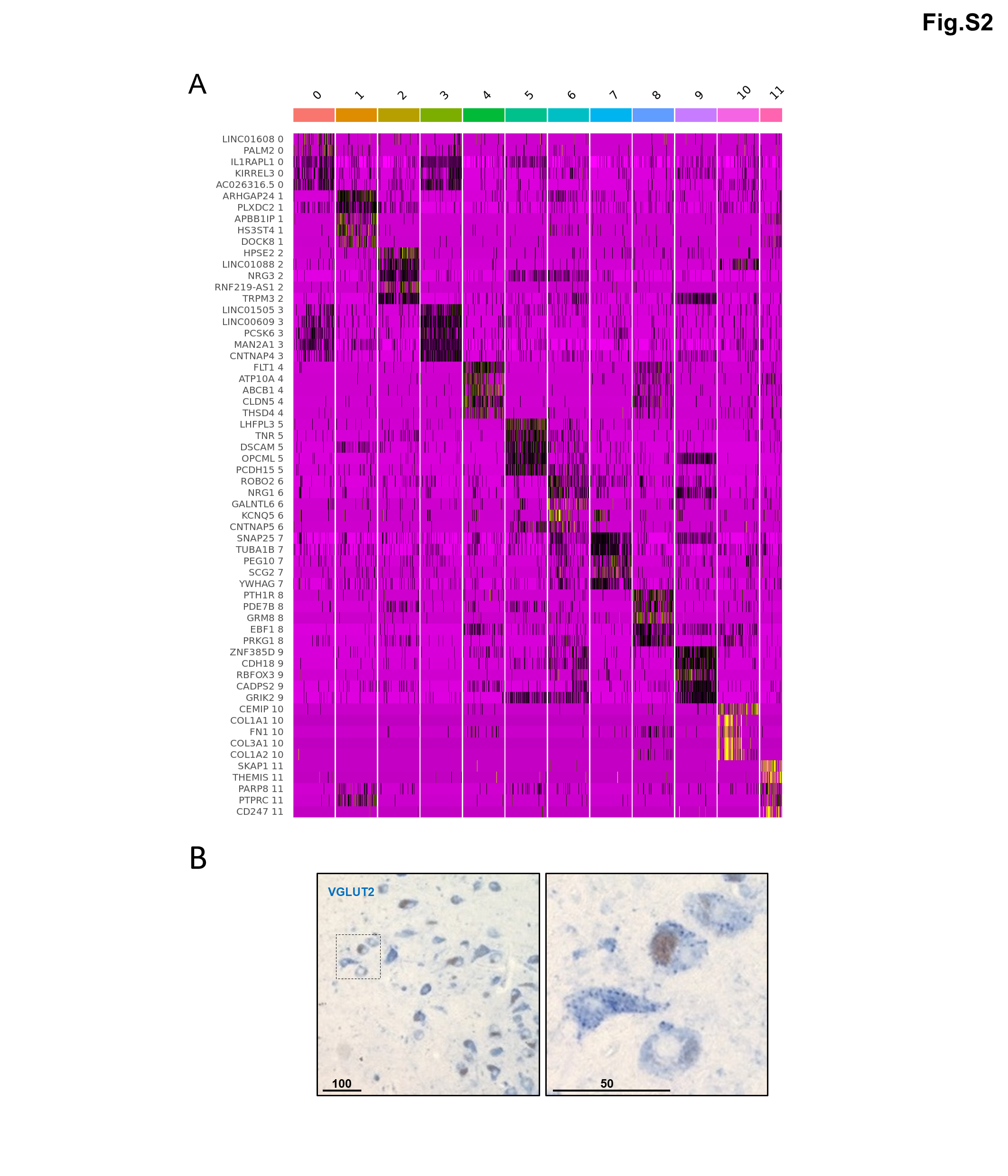

### FIG S3

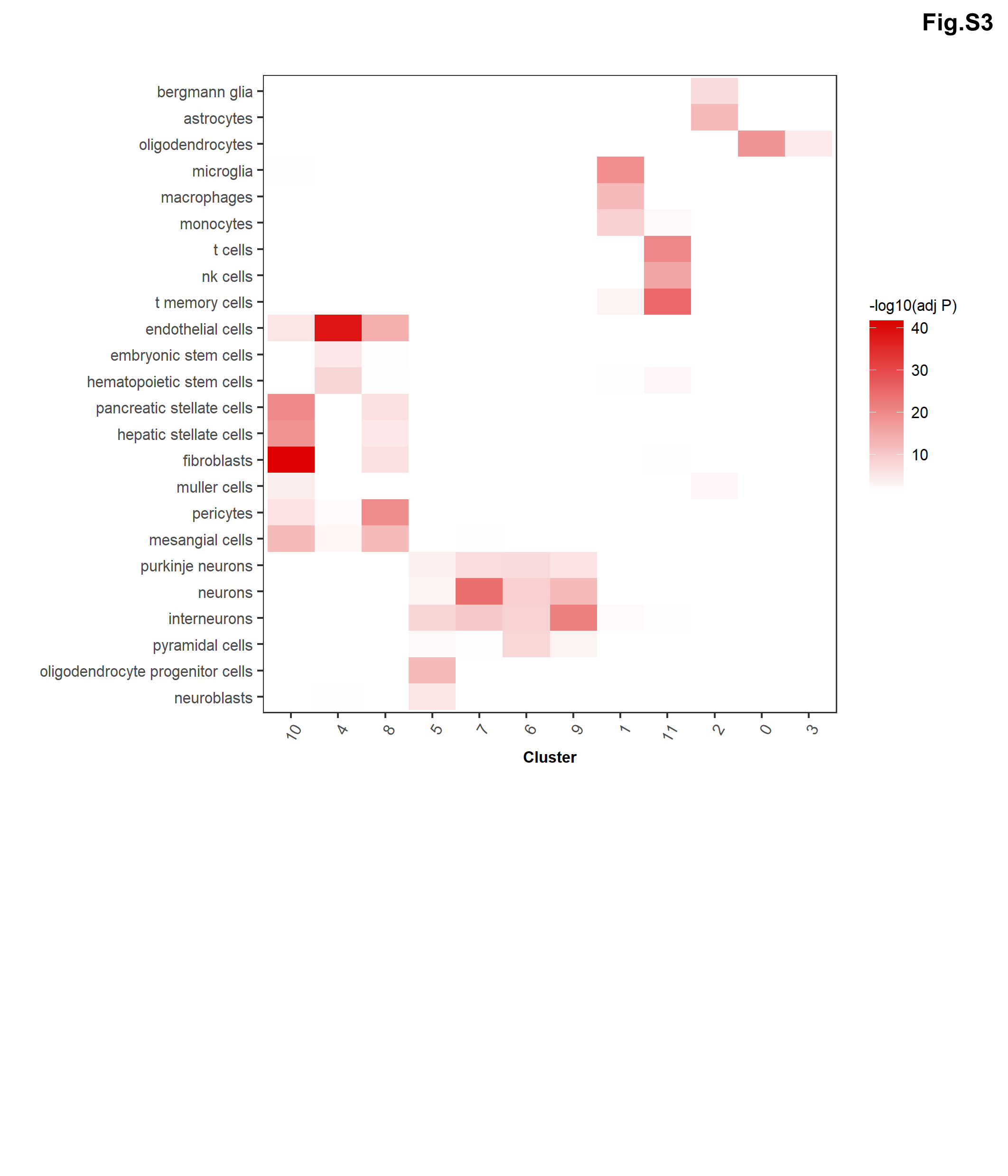

### FIG S4

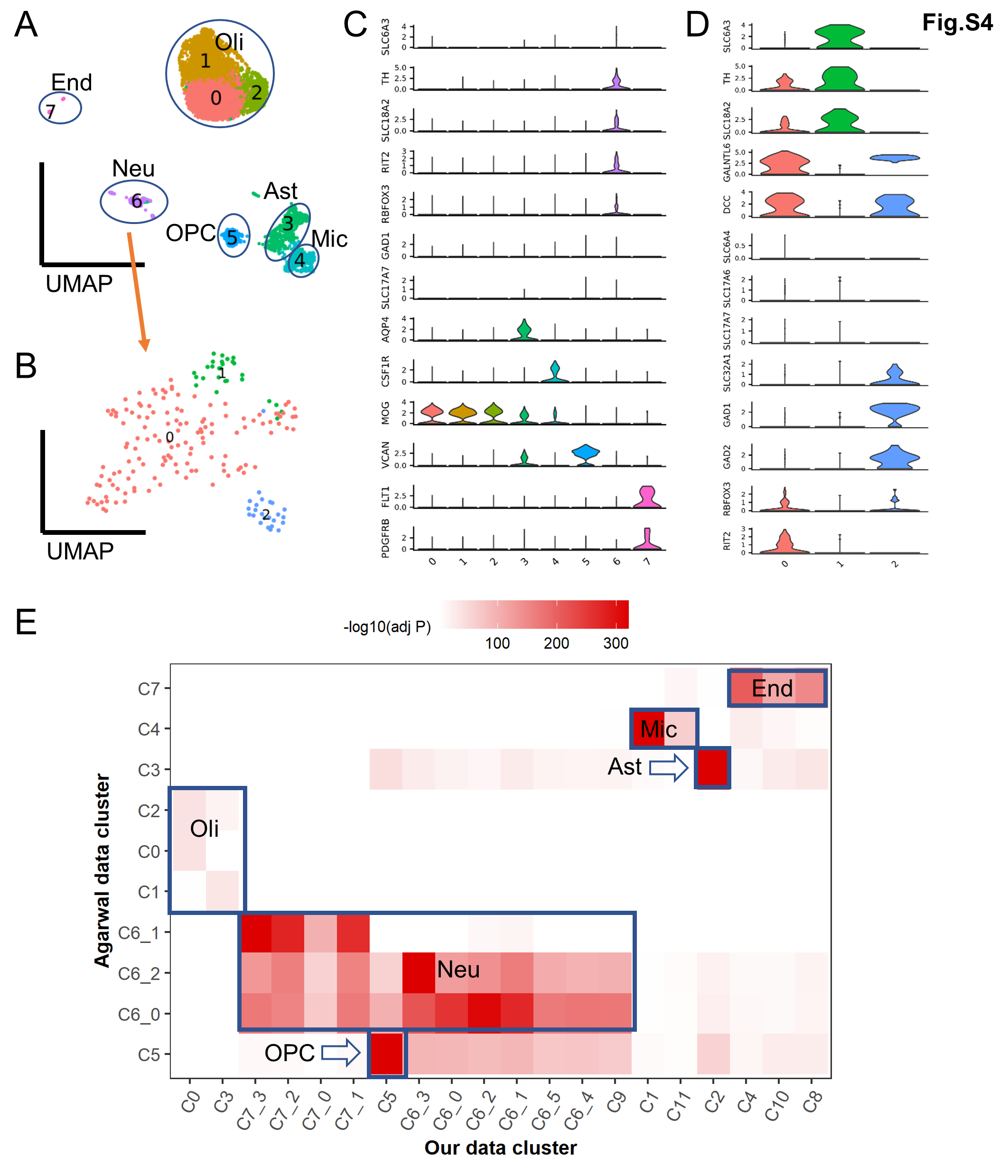

### FIG S5

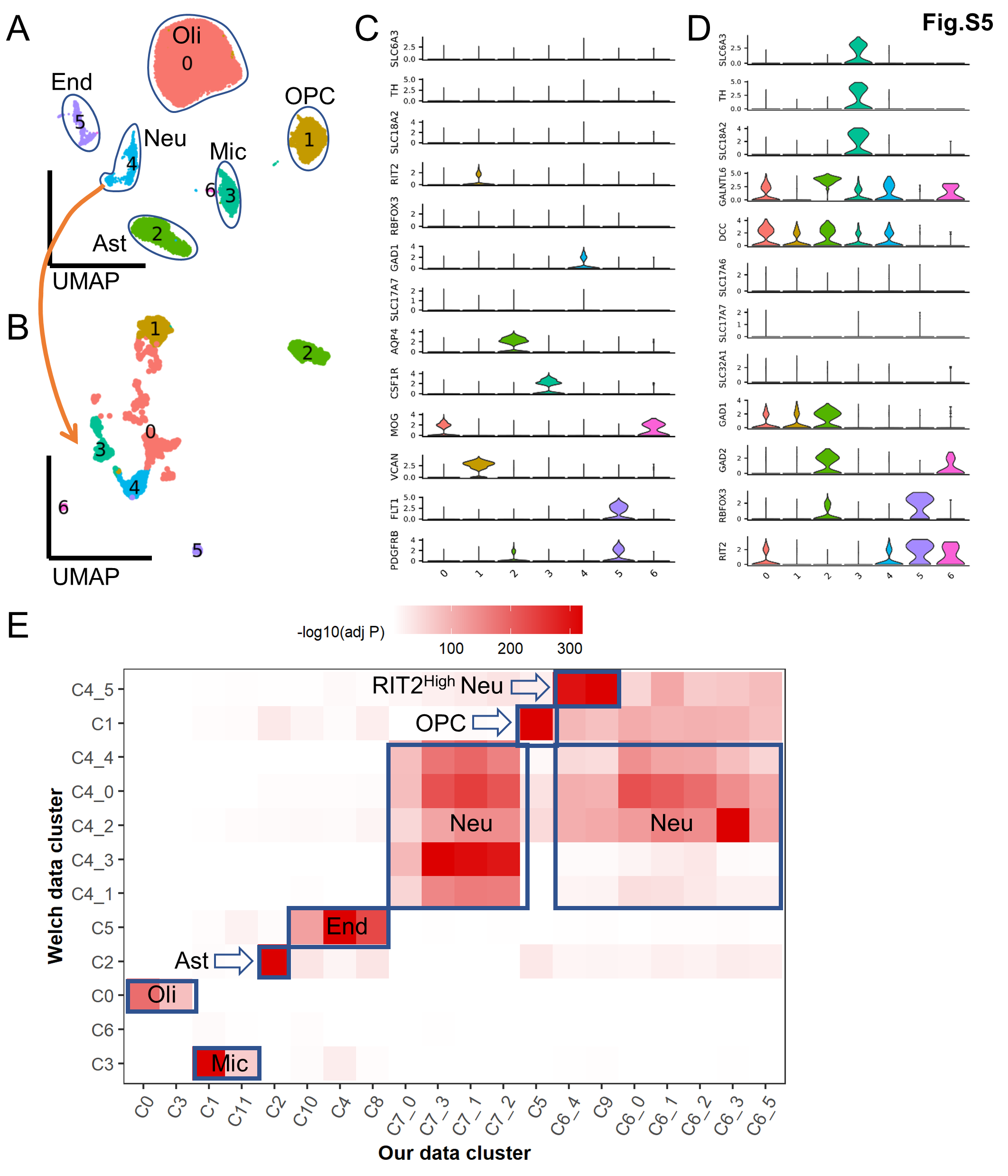

### FIG S6

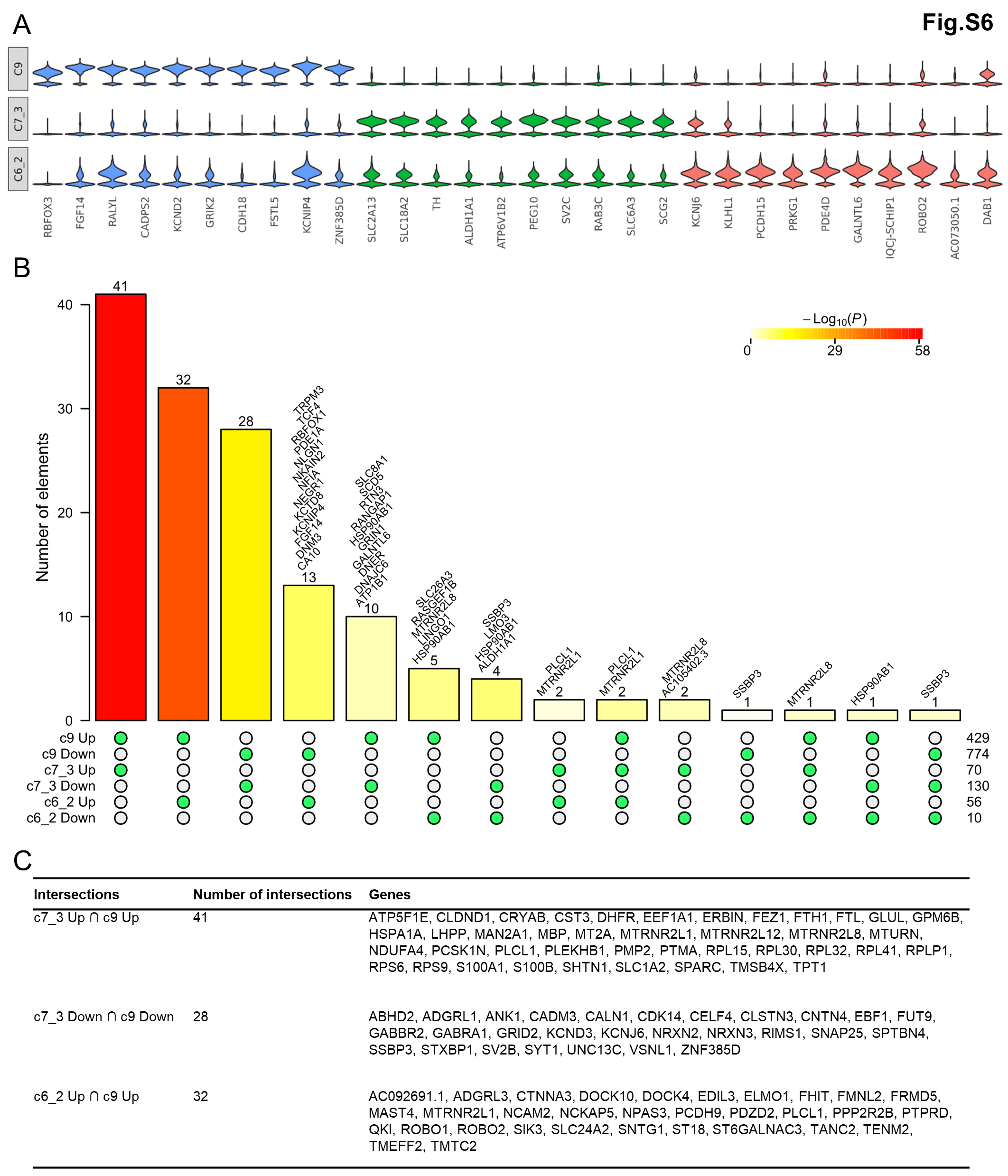

### FIG S7

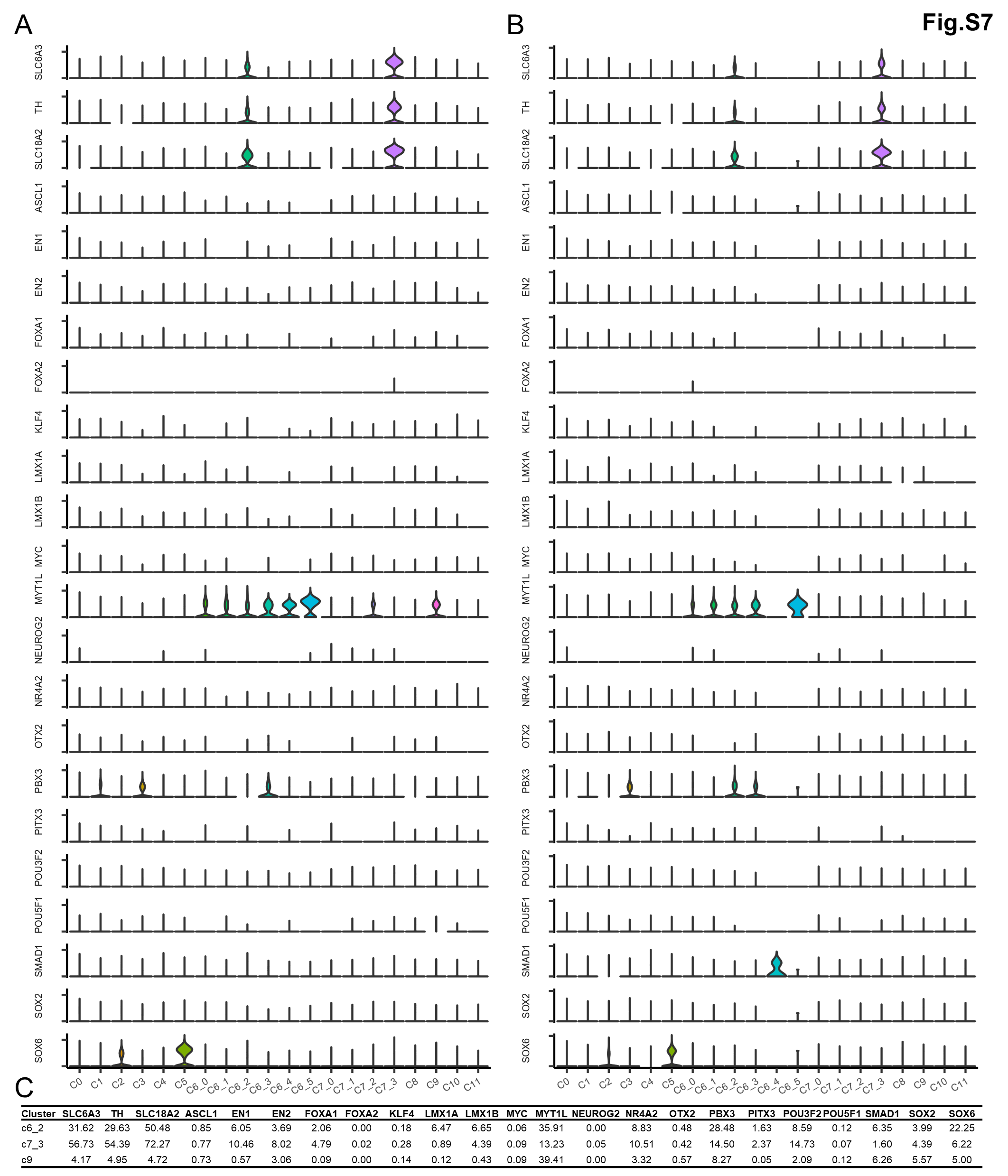

### FIG S8

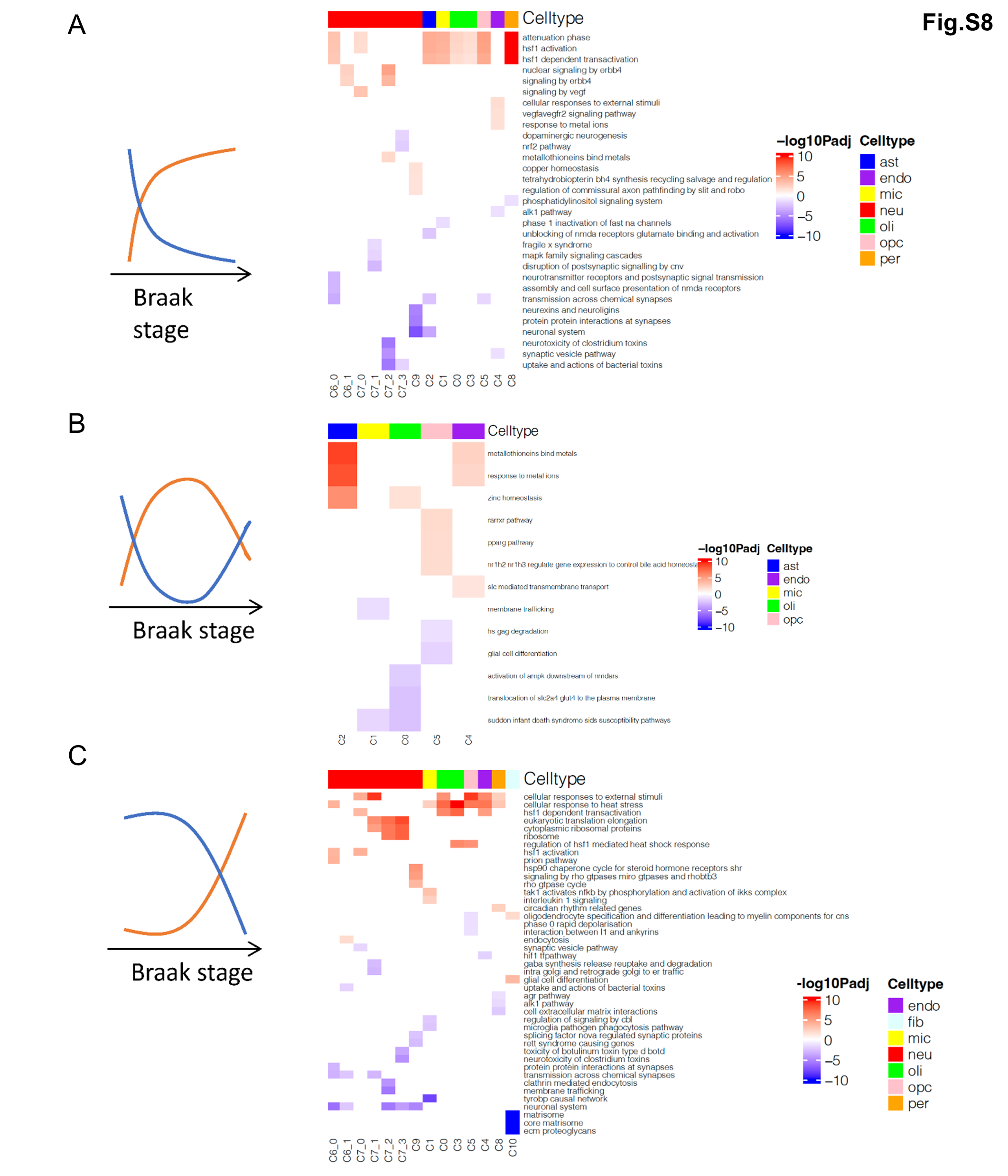

### FIG S9

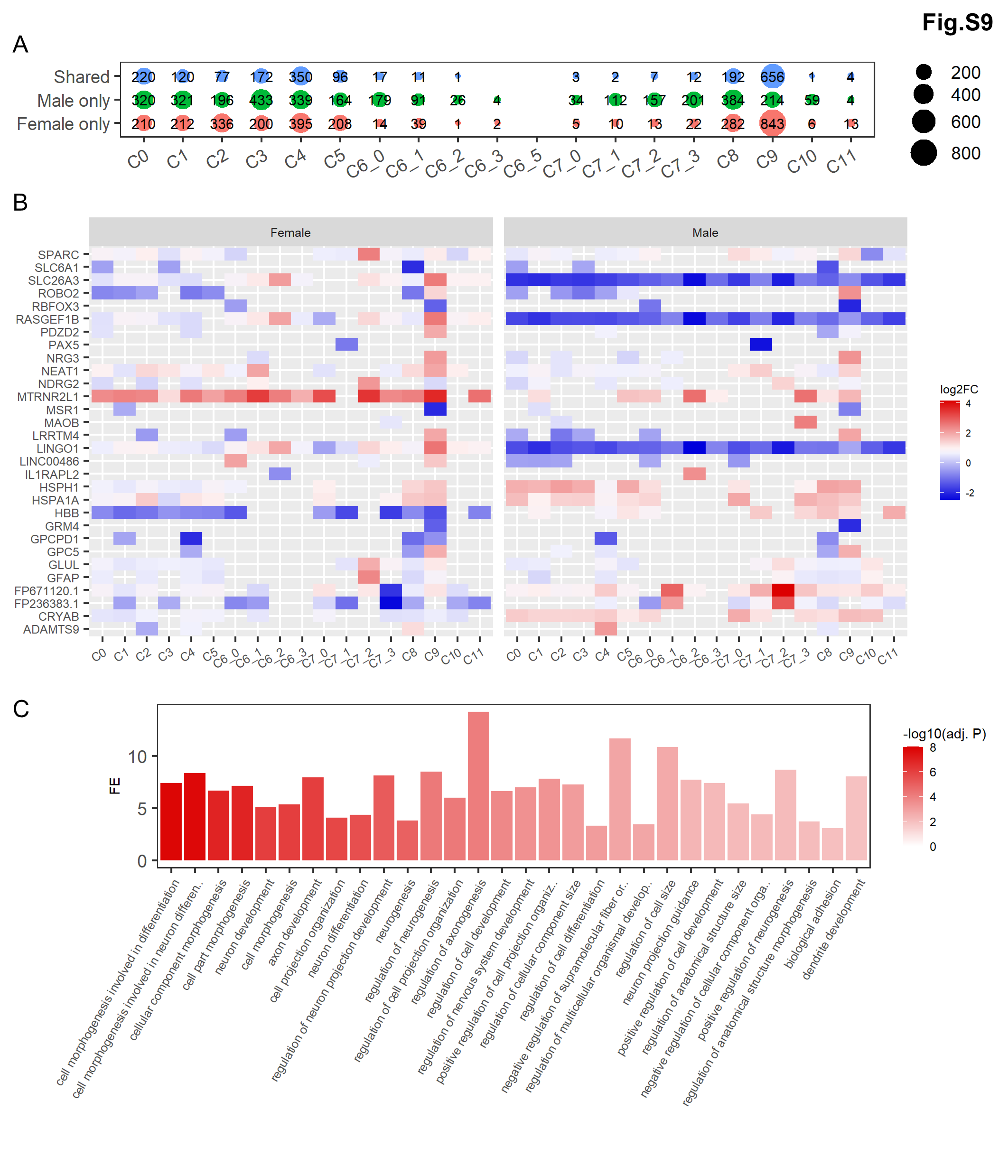

### FIG S10

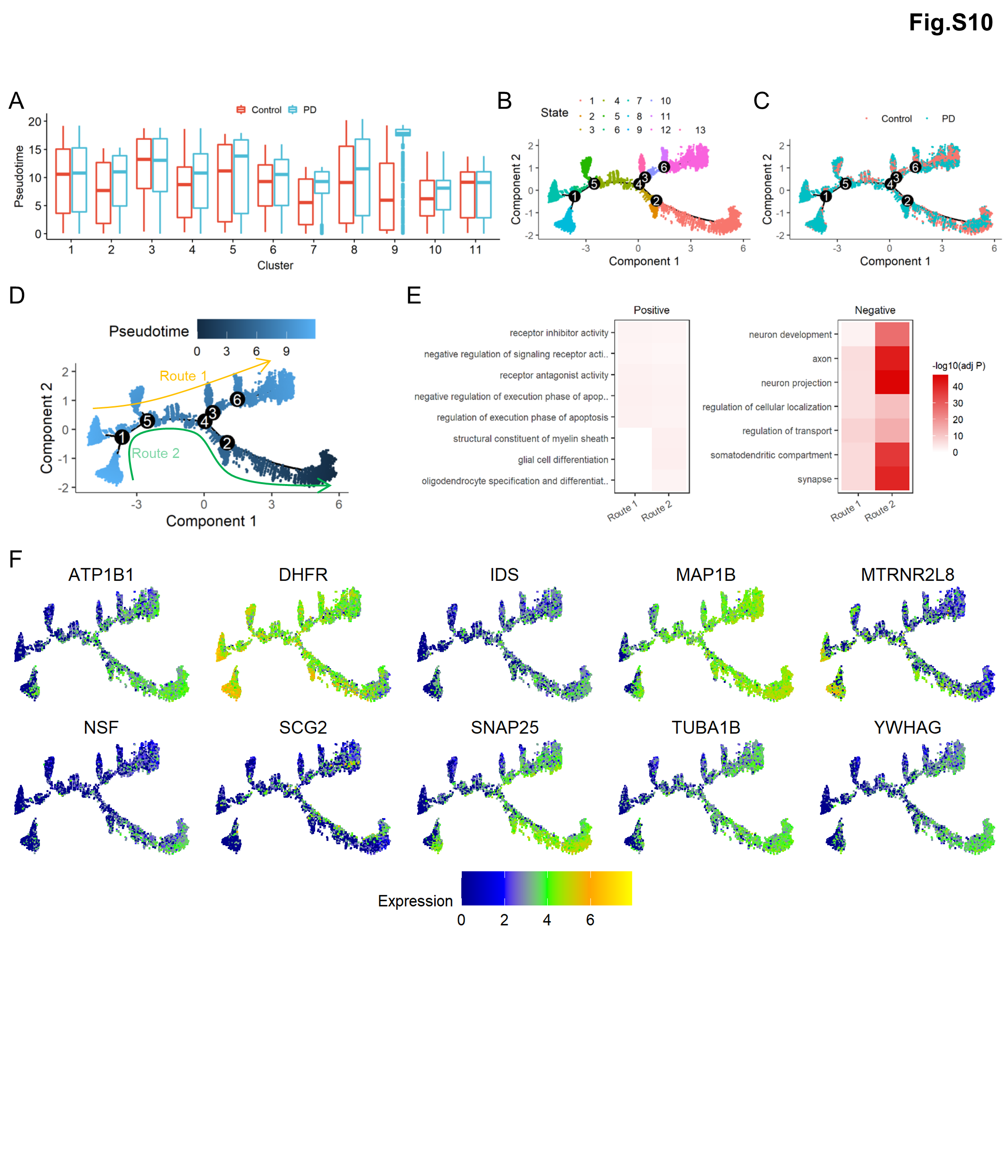
